# An ExbD Disordered Domain Peptide Inhibits TonB System Activity

**DOI:** 10.1101/2020.01.05.895219

**Authors:** Dale R. Kopp, Kathleen Postle

## Abstract

The TonB system energizes transport of essential nutrients, such as iron siderophores, across unenergized outer membranes of Gram-negative bacteria. The integral cytoplasmic membrane proteins of the TonB system--ExbB, ExbD, and TonB--transduce the protonmotive force of the cytoplasmic membrane to TonB-dependent outer membrane transporters for active transport. ExbD protein is anchored in the cytoplasmic membrane, with the majority of it occupying the periplasm. We previously identified a conserved motif within a periplasmic disordered domain that is essential for TonB system function. Here we demonstrated that export of a peptide derived from that motif into the periplasm prevented TonB system function and inhibited all known ExbD interactions in vivo. Formaldehyde crosslinking captured the ExbD peptide in multiple ExbD and TonB complexes. Furthermore, peptides with mutations in the conserved motif not only had significantly reduced ability to inhibit TonB system activity, but they also altered interactions with ExbD and TonB, indicating the specificity of the interaction. Conserved motif peptide interactions with ExbD and TonB mostly occurred between Stage II and Stage III of the TonB energy transduction cycle, a transition that is characterized by the use of protonmotive force. Taken together, the data suggest that the ExbD disordered domain motif has multiple interactions with TonB and ExbD during between Stage II and III of the TonB energization cycle. Because of the essentiality of the motif, it may be a potential template for design of novel antibiotics that target the TonB system.

**IMPORTANCE:** Gram-negative bacteria are intrinsically antibiotic-resistant due to the diffusion barrier posed by their outer membranes. The TonB system allows them to circumvent this barrier for their own nutritional needs, including iron. The ability of bacteria to acquire iron is a virulence factor for many Gram-negative pathogens. However, no antibiotics currently target the TonB system. Because TonB and ExbD must interact productively in the periplasm for transport across the outer membrane, they constitute attractive targets for potential antibiotic development where chemical characteristics need not accommodate the need to cross the hydrophobic cytoplasmic membrane. Here we show that a small ExbD-derived peptide can interfere with the TonB-ExbD interaction to inhibit the TonB system in vivo.

## INTRODUCTION

The TonB system is necessary under iron-limiting conditions for Gram-negative bacteria to acquire iron across their outer membranes. Because the TonB system is a virulence factor in many Gram-negative pathogens, it is an appealing target for novel antibiotics (1–5). Under iron-limiting conditions, such as in the human serum, Gram-negative bacteria secrete iron scavenging chelators called siderophores that bind iron with high affinity (6, 7). However, the iron-siderophores are too large, too scare, and too important to depend on diffusion through the outer-membrane porins, such as OmpC or OmpF, and require energy for active transport through larger 22-stranded beta-barrels called TonB dependent transporters (TBDTs) [recently reviewed in (8)]. Unlike the cytoplasmic membrane, there is not a sufficient energy source at the outer membrane such as a proton gradient or ATP molecules for active transport [reviewed in (9)]. Therefore, Gram-negative bacteria use the cytoplasmic membrane proteins of the TonB-system to transduce the energy from the proton gradient, called proton motive force (PMF), of the cytoplasmic membrane to the TonB-dependent transporters at the outer membrane (8).

The TonB system consists of various ligand-specific TonB-dependent transporters and three cytoplasmic membrane proteins ExbB, ExbD, and TonB. ExbB has three pass transmembrane domains (TMDs) with majority of the protein residing in the cytoplasm (10, 11).ExbD and TonB have a similar topology with a single pass TMD and a majority of the protein residing in the periplasm (12, 13). In Escherichia coli, the in vivo cellular ratio is 7 ExbB: 2 ExbD: 1 TonB (14), however the composition of a single active complex is unknown (14–18). ExbB serves as a scaffolding protein for both ExbD and TonB, and recently has been implicated in contributing to the signal transduction process (11, 19–22). ExbD and TonB transition through a series of interactions, categorized in stages (23), that describe how TonB energizes the TonB-dependent transporters. The TonB system must undergo a cyclic process in which TonB can energize and recycle to energize another TBDT because TonB is vastly outnumbered by the TBDTs (14, 21, 24). This process is referred to as the TonB energization cycle (21).

The ExbD and TonB interactions define the initial stages of the TonB energization cycle (23). Stage I is defined when no ExbD-TonB interaction is present. Stage II is defined by a PMF-independent ExbD-TonB interaction. Stage III is defined by a PMF-dependent ExbD-TonB interaction (23, 25). Although TonB is the only cytoplasmic membrane protein to directly contact the TBDTs, it is dependent on ExbD, ExbB, and PMF for function (19, 26, 27). ExbD has the only protonatable residue of the system within its transmembrane domain (residue D25) (11, 25, 28) and therefore has been hypothesized to be directly responsible for connecting PMF to TonB (25, 29). Because of its functional importance and periplasmic location, the ExbD could serve as a potential target for future antibiotics.

In *E. coli*, ExbD has a short C-terminal tail within the cytoplasm (residues ∼1-21), a single pass transmembrane domain (residues ∼22-43), and remainder of the protein in the periplasm (residues ∼44-141) (13). The periplasmic domain of ExbD is highly dynamic with multiple protein interfacial transitions (24, 28, 29, Kopp and Postle manuscript submitted). During signal transduction, residues within the ExbD periplasmic domain form many complexes – ExbD homodimers, two conformationally distinct ExbD-ExbB complexes, and PMF independent and dependent interactions (24, Kopp and Postle manuscript submitted). Different interfaces can involve the same ExbD residues which highlights the complexity of these interactions (29, Kopp and Postle, manuscript submitted).

Previous studies have characterized the ExbD protein in vivo. Any 10 amino acid deletion within the ExbD periplasmic domain rendered the protein non-functional. Deletions specifically within the 92-121 region inhibited all known ExbD complexes – ExbD-ExbB, ExbD-TonB, and ExbD homodimers (31). Further characterization within this region has identified functionally important residues involved in the PMF-dependent ExbD-TonB interaction (29). In a different study, single residues with ExbD disordered region (residues 44-63) could trap all known ExbD complexes via in vivo photo-cross-linking (Kopp and Postle, manuscript submitted). A conserved motif had been identified within the 44-63 region in which alanine substitutions within the motif prevented ExbD from transitioning to the PMF-dependent ExbD-TonB interaction from the PMF-independent ExbD-TonB interaction (Kopp and Postle, manuscript submitted).

Although there are many functional regions within the ExbD periplasmic domain to potentially target with a novel antibiotic, disrupting specific ExbD interactions has not been demonstrated. TonB exhibits a dominant negative gene-dosage effect (1) meaning that when TonB is overexpressed, it inhibits TonB system activity rather than enhancing it. Studies have shown that expressing the periplasmic domain of TonB (residues 33-239) and truncations of this domain (residues 103-239 and 122-239) also inhibit TonB system activity (2) in which the authors concluded the effect was due to inhibition of the TonB-TBDT interactions. However, because the TBDTs are highly abundant, sequentially different, and nutrient redundant, potentially targeting TBDT-TonB interactions with an antibiotic likely would not be ideal. On the other hand, targeting the periplasmic protein-protein interactions between ExbD and TonB would more fruitful because the number of potential antibiotic targets would be substantially less than those of the TonB-TBDT interactions, and a TonB-ExbD target would likely have an identical interaction, and therefore make for a more specific target. Although many TonB and ExbD protein-protein interactions have been identified, it has not been demonstrated that any could be directly inhibited, including the functionally important ExbD-TonB PMF-dependent interaction.

Here, we tested the ability of 20-30 amino-acid peptides derived from ExbD to inhibit TonB system function. Using an intracellular approach by fusing the ExbD peptides to the periplasmic localizing cleavable signal sequence of dsbA, we found that a peptide of the ExbD disordered region (residues 44-63) significantly inhibited TonB system activity. Moreover, the dsbA(ss)-ExbD(44-63) peptide inhibited the ExbD-TonB PMF-dependent interaction and was captured in multiple complexes with both ExbD and TonB proteins. The dsbA(ss)-ExbD (44-63) peptide interactions with native ExbD were influenced by the presence of TonB whereas the peptide interactions with native TonB were dependent on ExbD. However, neither the peptide interactions with ExbD nor TonB were dependent of PMF. In addition, alanine substitutions for residues V45 and V47 and a deletion of 44-48 within the inhibitory peptide significantly reduced its potency as an inhibitor. These data demonstrate that residues within the conserved motif of the ExbD (44-63) peptide are important for inhibiting TonB system activity. Taken together, ExbD residues 44-48 are the key residues of the inhibiting peptide and may serve as a template for designing an inhibitor against the TonB system.

## RESULTS

### Periplasmic domain of ExbD can inhibit TonB system activity and inhibit formation of ExbB-TonB and ExbD-TonB complexes

A previous study showed that the periplasmic domain of TonB (residues 33-239) inhibited TonB system activity, which was attributed to blockage of the TonB-TBDT interactions (33). However, at the time of the study, the importance of the periplasmic PMF-dependent ExbD-TonB interaction was not known (25). To test whether the TonB periplasmic domain could inhibit the periplasmic PMF-dependent ExbD-TonB interaction the formaldehyde cross-linked complex was analyzed in cells overexpressing the TonB periplasmic domain.

For periplasmic localization, a construct was engineered with the TonB periplasmic domain (TonB residues 33-239) fused to the cleavable signal peptide sequence of the outer membrane protease, ompT [ompT(ss)] (34) with a four amino acid linker sequence [ompT(ss)-TonB(33–239)] (Fig. 1A; Fig. S1). Consistent with previous findings, the cells overexpressing the periplasmic domain of TonB [ompT(ss)-TonB(33–239)] exhibited an iron-deficient growth deficiency and were incapable of ^55^Fe-ferrichrome transport (Fig. S2 and data not shown). To determine if overexpressing ompT(ss)-TonB(33–239) prevented the formation of the PMF-dependent ExbD-TonB complex, the formaldehyde cross-linked complexes from WT cells were compared with those from WT cells overexpressing the ompT(ss)-TonB(33–239). Due to the indiscriminate detection of the TonB protein and the ompT(ss)-TonB(33–239) peptide, and the non-specific background cross-linking from the peptide, anti-ExbD was used rather than anti-TonB to probe the PMF-dependent ExbD-TonB complex (Fig. S3A).

**Figure 1:**
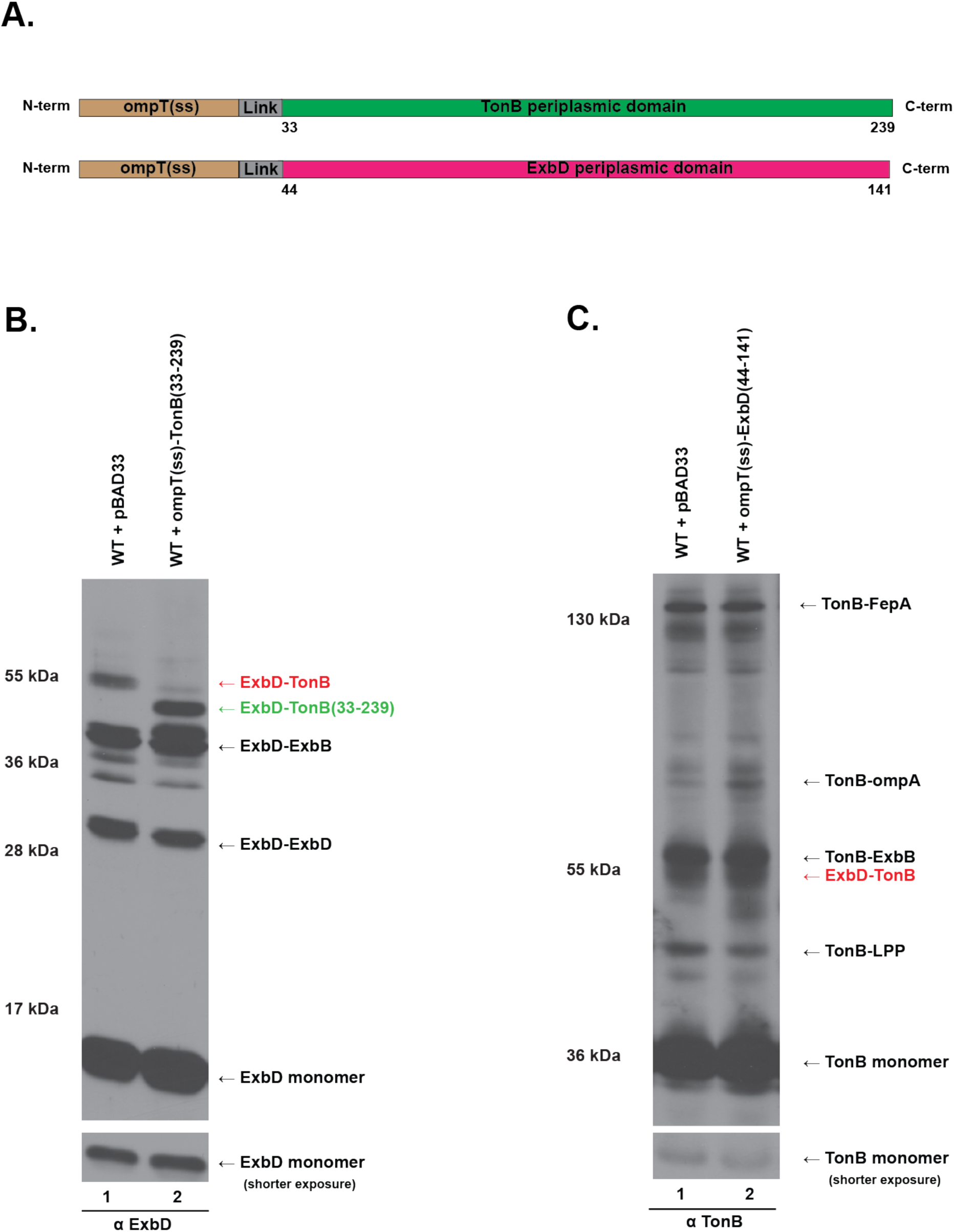
TonB and ExbD periplasmic domains inhibit the ExbD-TonB PMF-dependent interaction. (A) Schematic representations of the TonB (top) and the ExbD (bottom) periplasmic domains fused to the ompT signal sequences with linkers [ompT(ss)] from vector pET12a. The TonB periplasmic domain consisted of TonB residues 33-239 and the ExbD periplasmic domain consisted of ExbD residues 44-141. Linker sequences of 4 additional amino acids separated cleavage sites from the TonB and ExbD periplasmic domains (Fig. S1). (B) Formaldehyde cross-linking of W3110 (WT) expressing ompT(ss)-TonB(33-239) probed with anti-ExbD polyclonal antibodies. (lane 1) WT + pBAD33 refers to W3110 harboring the empty vector pBAD33. (lane 2) WT + ompT(ss)-TonB(33-239) refers to W3110 expressing plasmid-encoded ompT(ss)-TonB(33-239) under the arabinose inducible promoter (pKP1715). Positions of the previously characterized ExbD formaldehyde cross-linked complexes are shown: the PMF-dependent ExbD-TonB interaction (ExbD-TonB, red text), ExbD-ExbB heterodimer (ExbD-ExbB), and the ExbD homodimer (ExbD-ExbD) (25). ExbD-TonB(33-239) identifies ExbD complexed with TonB(33-239) (green text). (B, left) Mass markers are shown. (B, bottom) A shorter exposure of TonB monomer corresponding to each sample. The plasmid identities are listed in Table S1. (C) Formaldehyde cross-linking of W3110 (WT) expressing ompT(ss)-ExbD(44-141) probed with anti-TonB monoclonal antibodies. (lane 1) WT + pBAD33 refers to W3110 harboring the empty vector pBAD33. (lane 2) WT + ompT(ss)-ExbD(44-141) refers to W3110 expressing plasmid-encoded ompT(ss)-ExbD(44-141) under the arabinose inducible promoter (pKP1714). Positions of the previously characterized TonB formaldehyde cross-linked complexes are shown: TonB-FepA complex, the TonB-ExbB complex, the PMF-dependent TonB-ExbD complex (red text), TonB-LPP (Braun’s lipoprotein) and TonB monomer (25, 35). (C, left) Mass markers are shown. (C, bottom) A shorter exposure of ExbD monomer corresponding to each sample. The plasmid identities are listed in Table S1.

Cells overexpressing ompT(ss)-TonB(33–239) inhibited the PMF-dependent ExbD-TonB interaction (Fig. 1B). The ompT(ss)-TonB(33–239) expression however did not significantly prevent the formation of ExbD homodimers or ExbD-ExbB heterodimers, demonstrating that the ompT(ss)-TonB(33–239) peptide was specifically preventing the PMF-dependent ExbD-TonB interaction (Fig 1B). Since the PMF-dependent ExbD-TonB interaction occurs prior to TonB energetically interacting with the TBDTs (15), the loss of TonB system activity caused by the ompT(ss)-TonB(33–239) peptide may be from inhibiting the PMF-dependent ExbD-TonB interaction rather than the TonB-TBDTs interactions as previously proposed (33).

There is no current evidence that ExbD directly interacts with the TBDTs. Unlike TonB, ExbD neither formaldehyde cross-links with the TBDTs nor does it fractionate with the outer membrane (25, Ollis and Postle unpublished data). Therefore, to identify if the target of the ompT(ss)-TonB(33–239) peptide is disruption of PMF-dependent ExbD-TonB interaction or is disruption of the TonB-TBDT interaction, a construct of the ExbD periplasmic domain (ExbD residues 44-141) fused to ompT(ss) [ompT(ss)-ExbD(44-141)] was engineered and [Fig. 1A (bottom)]. Surprisingly, cells expressing this construct did not exhibit a noticeable iron-dependent growth deficiency and the PMF-dependent ExbD-TonB interaction was not inhibited (Fig. S2 and Fig. 1C). OmpT(ss) uses a posttranslational periplasmic delivery mechanism, and therefore it may not be ideal for quickly folding proteins (37), which may be the case with ExbD. To investigate the possibility of insufficient periplasmic localization of the ompT(ss)-ExbD (44-141) peptide, ExbD (44-141) was fused to the cleavable signal peptide sequence of dsbA [dsbA(ss)], which is known to use a co-translational periplasmic delivery mechanism (38, 39, Fig. S1, Fig. 2A).

**Figure 2:**
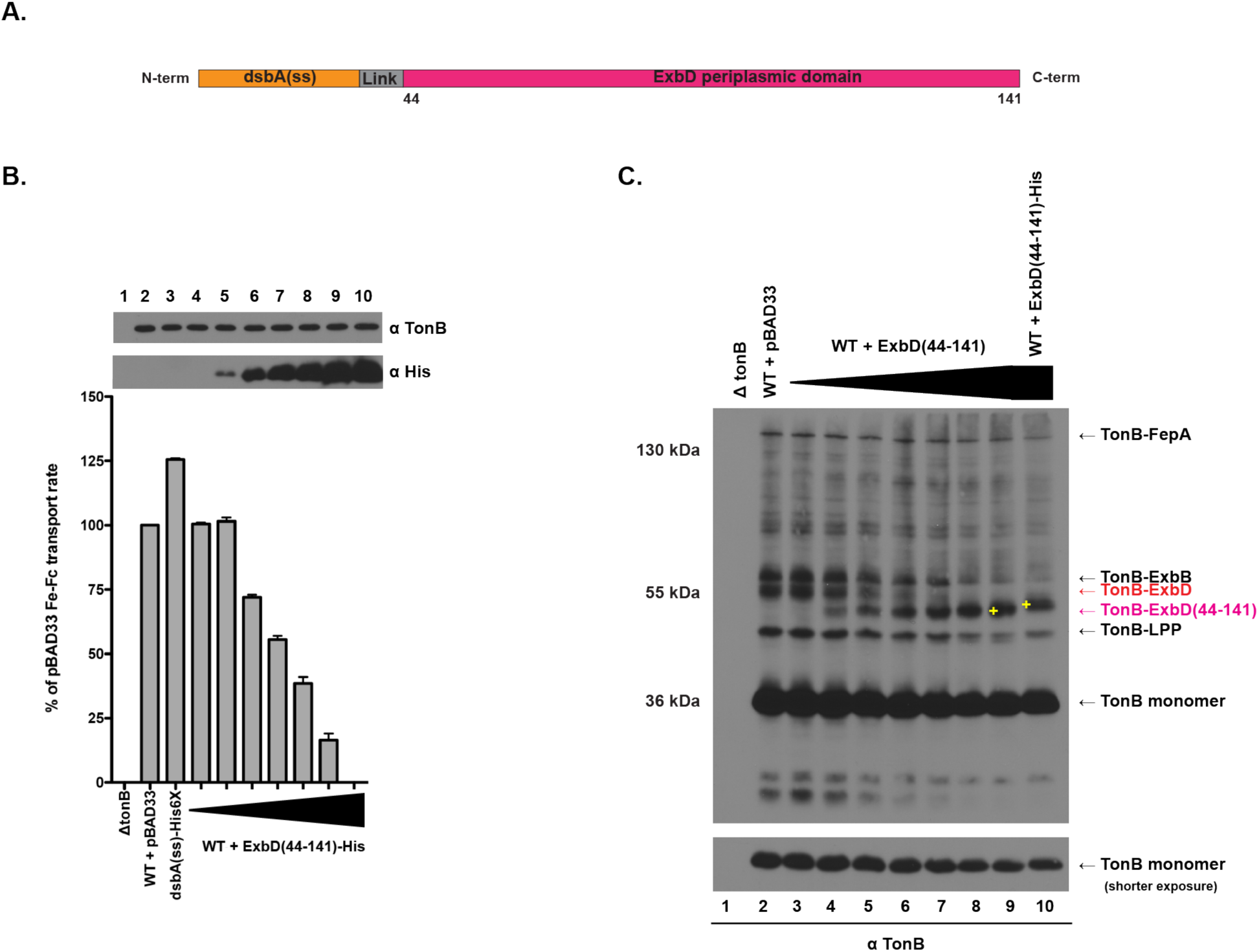
The ExbD periplasmic domain inhibits TonB system activity and prevents TonB-ExbB and TonB-ExbD complex formation. (A) A schematic representation of the ExbD periplasmic domain fused to dsbA signal sequence with a linker sequence [dsbA(ss)]. The ExbD periplasmic domain consists of ExbD residues (44-141). Linker sequences of 4 additional amino acids separated the dsbA signal peptidase cleavage site and the ExbD periplasmic domain (Fig. S1). (B) Initial rate of ^55^Fe-ferrichrome transport of W3110 (WT) with increasing expression of the ExbD periplasmic domain [ExbD(44-141)], and corresponding steady-state TonB protein and dsbA(ss)-ExbD(44-141)-His6X peptide levels. The ^55^Fe-ferrichrome transport rate was measured as described in the Materials and Methods section. (B, top) Steady-state protein expression levels of TonB and the dsbA(ss)-ExbD(44-141)-His6X peptide were probed with anti-TonB monoclonal antibodies and anti-His monoclonal antibodies, respectively. (B, left) The ^55^Fe-ferrichrome transport rate for each sample is recorded as the percent of the W3110 with pBAD33 (WT + pBAD33) transport rate. (lane 1) ΔtonB refers to W3110 with the tonB gene deleted (KP1477) and (lane 2) WT + pBAD33 refers to W3110 harboring the empty vector pBAD33. (lane 3) dsbA(ss)-His6X refers to W3110 expressing a plasmid-encoded dsbA signal sequence plus the four amino-acid linker sequence fused to C-terminal-His6X under the arabinose-inducible promoter (pKP1977). (lanes 4-10) ExbD(44-141)-His6X refers to W3110 expressing plasmid-encoded dsbA(ss)-ExbD(44-141)-His6X. The expanding black triangle represents the increasing arabinose that was added to each sample: 0%, 0.0025%, 0.005%, 0.01%, 0.02%, 0.05%, and 0.2% (w/v). (C) Formaldehyde cross-linking of W3110 (WT) with increasing expressions of dsbA(ss)-ExbD(44-141) probed with anti-TonB monoclonal antibodies. (lane 1) ΔtonB refers to W3110 with the *tonB* gene deleted (KP1477) and (lane 2) WT + pBAD33 refers to W3110 harboring the empty vector pBAD33. (lanes 3-9) WT + ExbD(44-141) is W3110 expressing plasmid-encoded dsbA(ss)-ExbD(44-141) under the arabinose inducible promoter (pKP1832). The expanding black triangle represents the increasing arabinose that was added to each sample: 0%, 0.0025%, 0.005%, 0.01%, 0.02%, 0.05%, and 0.2% (w/v). (lane 10) WT + ExbD(44-141)-His is W3110 expressing plasmid-encoded dsbA(ss)-ExbD(44-141)-His6X under the arabinose inducible promoter (pKP2013) with at 0.2% (w/v) arabinose. (C, right) The previously characterized TonB formaldehyde cross-linked complexes are shown: TonB-FepA complex, the TonB-ExbB complex, the PMF-dependent TonB-ExbD complex, TonB-LPP (Braun’s lipoprotein) and TonB monomer (25, 35). (C right, labeled in purple) The suspected complex of dsbA(ss)-ExbD(44-141) peptide trapped with TonB. The yellow plus sign identifies the mass shift between the suspected dsbA(ss)-ExbD(44-141)-TonB complex and dsbA(ss)-ExbD(44-141)-His6X-TonB complex. (C, left) Mass markers are shown. (C, bottom) A shorter exposure of TonB monomer corresponding to each sample. The plasmid identities are listed in Table S1.

To test whether expressing dsbA(ss)-ExbD(44-141) peptide was capable of inhibiting TonB system activity, the initial rate of TonB-dependent ^55^Fe-ferrichrome transport was measured in cells expressing this construct at increasing concentrations (40). A His-tag was fused to the C-terminus of the dsbA(ss)-ExbD(44-141) construct [dsbA(ss)-ExbD(44-141)-His6X] and was used to detect relative concentrations of the peptide via immunoblotting with an anti-His antibody. In a dose-dependent manner, expression of dsbA(ss)-ExbD(44-141)-His6X inhibited TonB-dependent ^55^Fe-ferrichrome transport (Fig. 2B). Cells with the highest expression of the dsbA(ss)-ExbD(44-141) were completely inactive. This demonstrated that the expression of the ExbD periplasmic domain, like that of TonB, completely inhibited TonB function. WT cells harboring the dsbA(ss)-His6X construct had a slight increase in ^55^Fe-ferrichrome transport rate for unknown reasons. However, because this construct did not inhibit TonB activity, the interpretation of the results was not affected.

To test whether the dsbA(ss)-ExbD(44-141) could prevent the formation of the PMF-dependent ExbD-TonB interaction, the WT formaldehyde cross-linked complexes were compared to those from WT expressing various concentrations of the dsbA(ss)-ExbD(44-141) peptide. In a dose-dependent manner, the dsbA(ss)-ExbD(44-141) prevented the PMF-dependent ExbD-TonB complex (Fig 2C). In addition to preventing the PMF-dependent ExbD-TonB interaction, and in contrast to the effect of the periplasmic TonB peptide on ExbD complexes, the dsbA(ss)-ExbD(44-141) peptide also prevented the TonB-ExbB complex. The PMF-dependent ExbD-TonB complex inhibition was the most sensitive to the dsbA(ss)-ExbD(44-141) peptide but interestingly, the ExbB-TonB interaction was also significantly prevented in cells expressing the dsbA(ss)-ExbD(44-141) peptide for reasons unknown. It is possible that this result suggests that ExbD and ExbB share a periplasmic interface with TonB. Alternatively, the ExbD periplasmic domain may interact with TonB in such a way that prevents TonB from assuming a conformation that is required for it to interact with ExbB.

Although the TonB-FepA and TonB-LPP complexes were slightly reduced in cells expressing dsbA(ss)-ExbD (44-141), the reduction did not appear to be significantly different from protein levels of TonB monomer (Fig. 2C, lane 9 and lane 2). Therefore, it is likely that the TonB-FepA and TonB-LPP complexes appeared reduced because of a lesser amount of TonB.

A ∼ 50 kDa unknown TonB complex was also captured in cells expressing the dsbA(ss)-ExbD(44-141) peptide. Considering its molecular weight and the fact that this complex increased with dsbA(ss)-ExbD(44-141) expression, it was likely the ∼ 50 kDa unknown TonB complex was TonB with the ExbD(44-141) peptide. To confirm the identity of this complex, this complex was compared to cells expressing the dsbA(ss)-ExbD(44-141)-His6X peptide. As expected, the unknown complex migrated slightly slower on the SDS-Page gel from cells expressing the dsbA(ss)-ExbD(44-141)-His6X peptide compared to those expressing the dsbA(ss)-ExbD(44-141) peptide. Therefore, this complex was identified as a TonB-ExbD(44-141) peptide complex.

### The ExbD disordered domain peptide (residues 44-63) inhibits TonB system activity

To identify the segments within dsbA(ss)-ExbD(44-141) peptide that were responsible for inhibiting TonB system activity, overlapping ExbD peptides of 20 – 30 amino acids that covered the entire ExbD periplasmic domain were fused to dsbA(ss) followed by a His6X (Fig. 3A) and tested for their ability to disrupt TonB system activity. Unfortunately, most of the dsbA(ss) ExbD peptides were proteolytically unstable and therefore, they could not be completely ruled as potential TonB system activity inhibitors. However, one peptide which corresponded to the ExbD disordered domain, dsbA(ss)-ExbD(44-63)-His6X, significantly inhibited the TonB-dependent ^55^Fe-ferrichrome transport rate (Fig. 3B).

**Figure 3:**
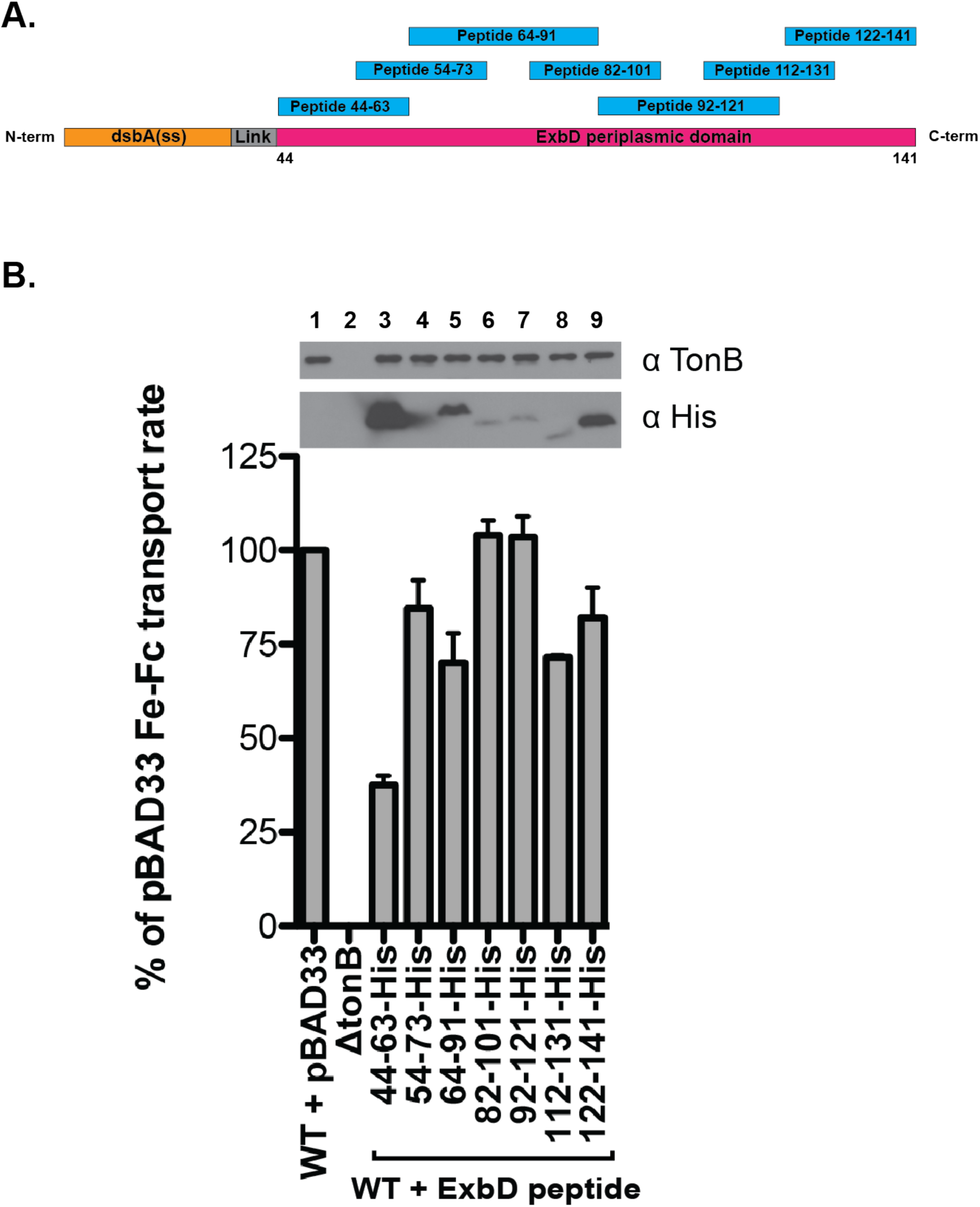
dsbA(ss)-ExbD(44-63)-His is the only peptide stable and inhibits TonB system activity. (A) A schematic representation of the dsbA(ss)-ExbD periplasmic domain with the overlapping ExbD peptide constructs. Overlapping peptides of 20-30 amino acids derived from the ExbD sequence spanning the periplasmic domain were fused to dsbA signal sequences [dsbA(ss)] with a four amino acid linker sequence. (B) Initial rate of ^55^Fe-ferrichrome transport of W3110 (WT) expressing peptides derived of the ExbD protein. The ^55^Fe-ferrichrome transport rate was measured as described in the Materials and Methods section. (B, top) Steady-state protein expression levels of TonB and the His-tagged peptides that were probed with anti-TonB monoclonal antibodies and anti-His monoclonal antibodies, respectively. (B, left) The ^55^Fe-ferrichrome transport rate was determined as the percent of the W3110 with the empty vector, pBAD33 (WT + pBAD33) transport rate. (lane 1) WT + pBAD33 refers to W3110 harboring the empty vector pBAD33, and (lane 2) ΔtonB refers to W3110 with the *tonB* gene deleted (KP1477). (lanes 3-9) 44-63-His, 54-73-His, 64-94-His, 82-101-His, 92-121-His, 112-131-His, and 122-141-His refer to W3110 expressing the plasmid-encoded dsbA(ss)-ExbD(44-63)-His6X, dsbA(ss)-ExbD(54-73)-His6X, dsbA(ss)-ExbD(64-91)-His6X, dsbA(ss)-ExbD(82-101)-His6X, dsbA(ss)-ExbD(92-121)-His6X, dsbA(ss)-ExbD(112-131)-His6X, and dsbA(ss)-ExbD(122-141)-His6X, respectively, under the arabinose-inducible promoter with 0.2% arabinose (w/v). The plasmid identities are listed in Table S1.

### The ExbD disordered domain peptide prevents ExbD and TonB formaldehyde cross-linked complexes and was captured with ExbD and TonB

Previously, the ExbD periplasmic disordered domain had been shown to be essential for forming the PMF-dependent ExbD-TonB interaction (31, Kopp and Postle manuscript submitted), and the dsbA(ss)-ExbD(44–63)-His6X fall within the ExbD periplasmic disordered domain. Here, we tested whether the expression of dsbA(ss)-ExbD(44-63) peptide prevents the PMF-dependent ExbD-TonB interaction.

Similar to cells that had expressed the dsbA(ss)-ExbD(44-141) peptide, cells expressing the dsbA(ss)-ExbD(44-63) peptide, were prevented from forming the PMF-dependent ExbD-TonB formaldehyde cross-linked complex (Fig. 4A). Furthermore, cells expressing this construct had reduced ExbB-ExbD heterodimers and ExbD homodimers compared to WT expressing the empty vector. Instead, cells expressing the dsbA(ss)-ExbD(44-63) peptide captured three unknown complexes with molecular weights of 17.1 kDa, 20.3 kDa and, 24.1 kDa (Fig 4A). To test whether any of these unknown ExbD complexes consisted of the dsbA(ss)-ExbD(44-63) peptide, the complexes were compared with those from cells that had expressed the dsbA(ss)-ExbD(44-63)-His6X peptide (Fig. 4A). The 20.3 kDa and, 24.1 kDa unknown ExbD complexes appeared to migrate slower on the SDS PAGE gel from the cells expressing the His6X tagged version of the peptide than from those expressing the peptide *without* the His6X tag. Because one of the complexes migrated twice as slow as the other (Fig. S5A), these complexes were identified as dsbA(ss)-ExbD(44-63) – ExbD and two-dsbA(ss)-ExbD(44-63) – ExbD (referred to as ExbD-peptide_1_ and ExbD-peptide_2_). However, the 17.1 kDa unknown complex that formed immediately above the ExbD monomer did not appear to migrate slower from cells expressing the His6X-tagged peptide(Fig. S5A). Therefore, the 17.1 kDa complex was not identified as a dsbA(ss)-ExbD(44-63) peptide-ExbD complex. Since the molecular weight of this complex was only approximately 1.5 kDa larger than ExbD monomer, it may be a conformational difference in ExbD. Although the 17.1 kDa complex was not identified as an ExbD-peptide complex, because it was only present in the dsbA(ss)-ExbD(44-63) peptide-expressing cells, it suggests that the peptide influences this conformation formation.

**Figure 4:**
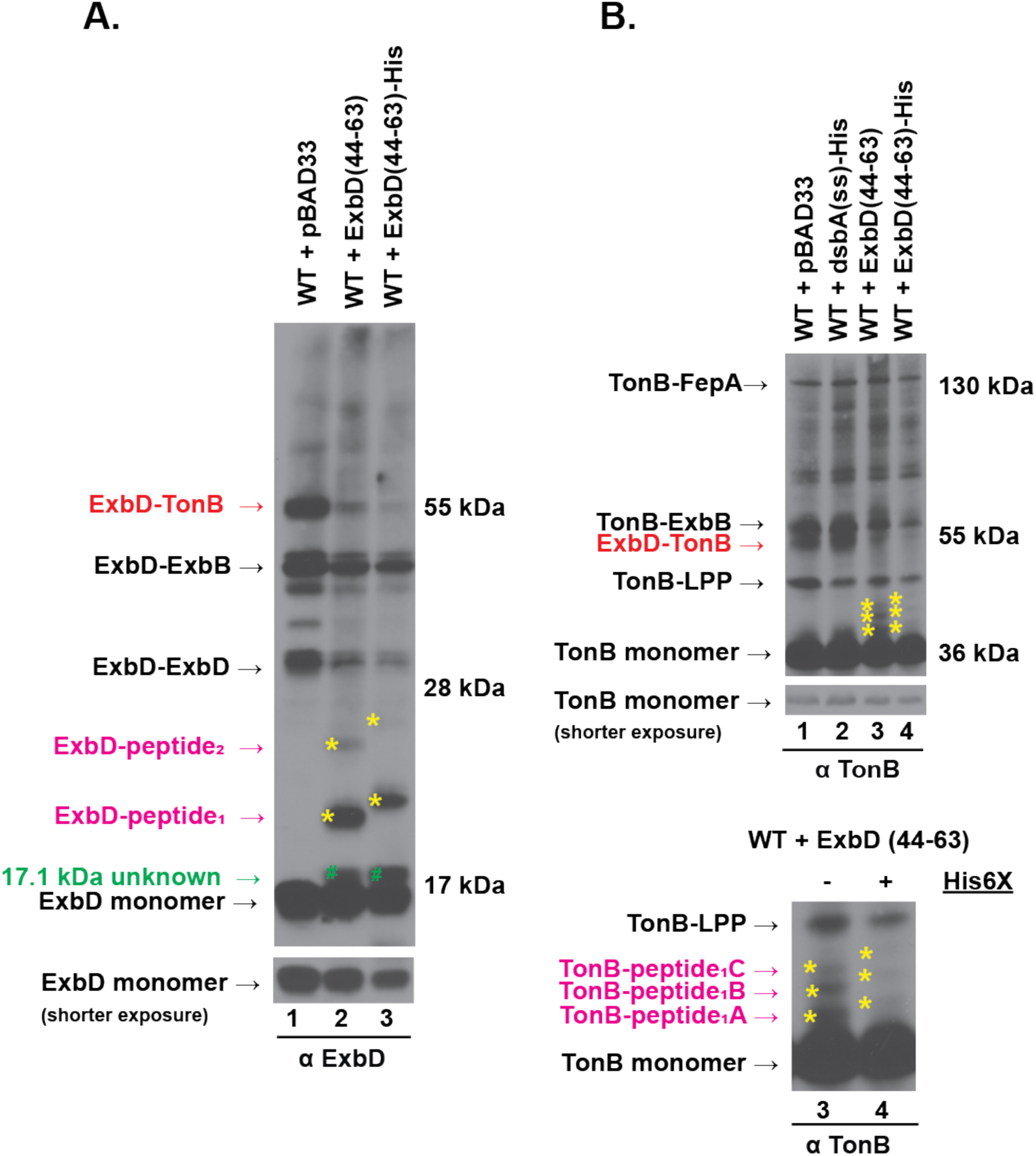
dsbA(ss)-ExbD(44-63) inhibits the TonB system activity, interacts with ExbD and TonB and inhibits the ExbD-TonB complex. (A) Formaldehyde cross-linking of W3110 (WT) with a plasmid expressing of dsbA(ss)-ExbD(44-63) probed with anti-ExbD polyclonal antibodies. (lane 1) WT+ pBAD33 refers to W3110 harboring the empty vector pBAD33. (lane 2) WT + ExbD(44-63) is W3110 expressing the plasmid-encoded dsbA(ss)-ExbD(44-63) under the arabinose inducible promoter (pKP1847) with 0.2% (w/v) arabinose. (lane 3) WT + ExbD(44-63)-His refers to W3110 expressing the plasmid-encoded dsbA(ss)-ExbD(44-63)-His6X under the arabinose inducible promoter (pKP1960) with 0.2% (w/v) arabinose. (A, left) Positions of the previously characterized ExbD formaldehyde cross-linked complexes are shown: the PMF-dependent ExbD-TonB complex (red), the ExbD-ExbB complex, the ExbD homodimer (ExbD-ExbD), and ExbD monomer (25). (A, left, labeled in purple) The suspected complexes of the dsbA(ss)-ExbD(44-63) peptide trapped with ExbD. The yellow plus sign identifies the shift between dsbA(ss)-ExbD(44-63)-ExbD complex and the dsbA(ss)-ExbD(44-63)-His6X-ExbD complex. (A, right) Mass markers are shown. (A, bottom) A shorter exposure of ExbD monomer corresponding to each sample. Unidentified 17.1 kDa complex is labeled with a green pound sign. (B) Formaldehyde cross-linking of W3110 (WT) with a plasmid expressing of dsbA(ss)-ExbD(44-63) probed with anti-TonB monoclonal antibodies. Same samples from the same experiment as in (B) probed with anti-TonB monoclonal antibodies. (Top panel, left) Positions of the previously characterized TonB formaldehyde cross-linked complexes are shown TonB-FepA complex, the TonB-ExbB complex, the PMF-dependent TonB-ExbD complex, TonB-LPP (Braun’s lipoprotein) and TonB monomer (35, 25). (Top panel, right) Mass markers are shown. (Top panel, below) A shorter exposure of TonB monomer corresponding to each sample. (Bottom panel) A magnified image of the Top panel lanes 3 and 4 to resolve the suspected complexes of the dsbA(ss)-ExbD(44-63) peptide and those of dsbA(ss)-ExbD(44-63)-His6x peptide trapped with TonB (yellow asterisk). Migration difference between each corresponding complex trapped from expressing dsbA(ss)-ExbD(44-63) and that from expressing dsbA(ss)-ExbD(44-63)-His6X each shifted with identically. (Bottom panel, left) The suspected complexes of dsbA(ss)-ExbD(44-63) peptide trapped with TonB. The yellow asterisk identifies the dsbA(ss)-ExbD(44-63)-TonB complex shift without and with the His6X-tag. The plasmid identities are listed in Table S1.

From the TonB complex perspective, similar to the dsbA(ss)-ExbD(44-141) peptide expression, the dsbA(ss)-ExbD(44-63) expression also prevented cells from forming TonB-ExbB formaldehyde cross-linked complex in addition to that of the PMF-dependent ExbD-TonB (Fig. 4B). Like the effect on ExbD, the dsbA(ss)-ExbD(44-63) expression caused three unknown TonB complexes to form. All of these complexes migrated slower from cells expressing dsbA(ss)-ExbD(44-63)-His6X compared to those expressing dsbA(ss)-ExbD(44-63) (Fig. 4B). Therefore, all three unknown TonB complexes likely contained the peptide. However, the distances between each of three complexes was similar in cells expressing the His-tagged version of the peptide relative to cells expressing the peptide version *without* the His-tag (Fig. S5B), which suggests that each of these complexes did not vary in the number of peptides that were captured (referred to as TonB-peptide_1_A, TonB-peptide_1_B, and TonB-peptide_1_C). Although the reason(s) are unclear why there are molecular weight differences between these complexes, it may be that one peptide interacts with TonB in three different ways. This would be consistent with a previous finding that showed single photo-cross-linkable residues within the ExbD disordered domain captured multiple conformations of the ExbD-TonB interaction (Kopp and Postle, manuscript submitted).

### The periplasmic fraction of the ExbD disordered domain peptide is low

The dsbA(ss) uses a co-translational translocation mechanism to localize recombinant proteins to the periplasm (39). In co-translation, the translation of a dsbA(ss) construct is arrested after the signal sequence translation, which is recognized by a signal recognition particle (SRP). The ribosome-SPR-bound nascent peptide complex is delivered to the cytoplasmic membrane receptor, FtsY, and subsequently transferred to the SecYEG translocase for export (39). To determine the proportion of the dsbA(ss)-ExbD(44-63) peptide localized to the periplasm, proteinase K (PK) accessibly was tested on whole cells, spheroplasts, lysed spheroplasts, and the periplasm from cells expressing the peptide (Fig. 5). Whole cells have intact outer and cytoplasmic membranes and thus, proteins and peptides that are compartmentalized within the *E. coli* cells would be protected from PK degradation. Spheroplasts have intact cytoplasmic membranes, but they do not intact outer membranes. In these samples, the periplasmic fraction is removed and only the proteins compartmentalized within the cytoplasm would be protected from PK degradation. Lysed spheroplasts have no intact membranes, and therefore, no protein or peptide would be protected from PK degradation.

**Figure 5:**
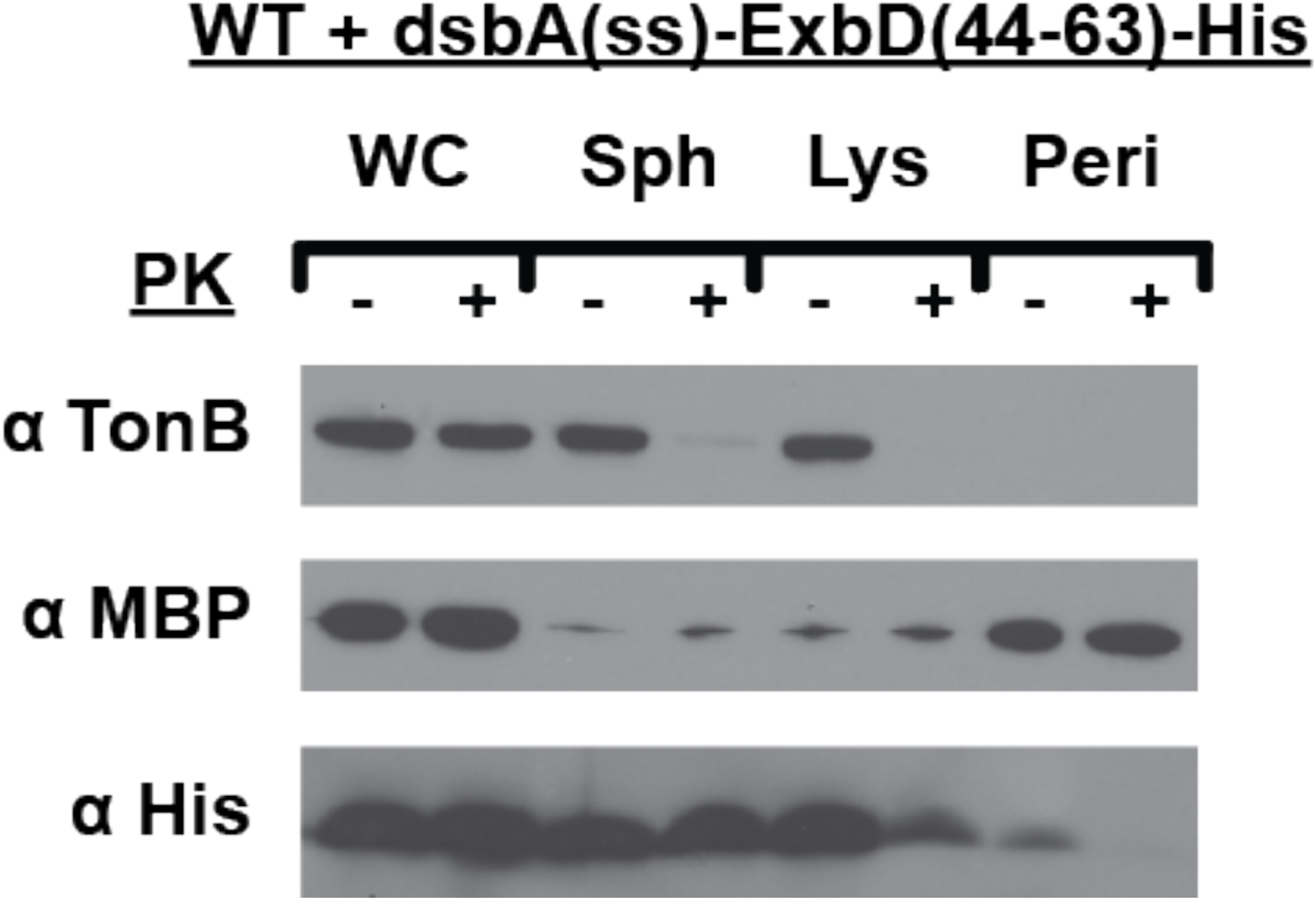
dsbA(ss)-ExbD(44-63) is inefficiently localized to the periplasm. Proteinase K (PK) accessibility in Whole cells (WC), Spheroplasts (Sph), Lysed Spheroplasts (Lys), and periplasmic fraction (Peri) of W3110 expressing plasmid-encoded dsbA(ss)-ExbD(44-63)-His6X (pKP1960). Strains were grown to an A_550_ of 0.2 at which point 0.2% (w/v) of arabinose was added to induce intracellular expression of the dsbA(ss)-ExbD(44-63)-His6X. Strains were grown until mid-exponential phase. WC, Sph, Lys, and Peri fractions treated (+) and untreated (-) with Proteinase K as described in the Materials and Methods. Equivalent numbers of cells were visualized on immunoblots of 16% acrylamide/ 6% bis-acrylamide SDS tricine gels. The periplasmic soluble maltose-binding protein (MBP) was probed with anti-maltose-binding protein monoclonal antibodies (Top panel). The cytoplasmic membrane-embedded, periplasmic TonB protein was probed with anti-TonB monoclonal antibodies (Bottom panel) and ExbD(44-63)-His peptide was probed with anti-His monoclonal antibodies. The MBP, TonB, and ExbD(44-63)-His were all probed from same immunoblot.

The bulk of TonB occupies the periplasmic space, rendering it accessible to PK in spheroplasts (54). However, because it is also tethered to the cytoplasmic membrane, it would not have been collected from the soluble periplasmic fraction. As expected, TonB was present in the whole cells, spheroplasts, and lysed spheroplasts but absent in the soluble periplasmic fraction. It was degraded by PK in the spheroplasts and lysed spheroplasts but not in whole cells. In contrast, the maltose-binding (MBP) is a soluble periplasmic protein that is not tethered to any membrane. As expected, MBP was predominantly present in the whole cells and periplasmic fractions. Interestingly, MBP was not degraded by PK in the periplasmic fraction, which suggests that it may have intrinsic resistance to PK. The TonB, and MBP were controls to verify the localization. Surprisingly, the soluble periplasm had only a small fraction of the dsbA(ss)-ExbD(44-63)-His6X peptide. The peptide was mostly degraded in the lysed spheroplasts, which suggests that most of the dsbA(ss)-ExbD(44-63)-His6X was localized to the cytoplasm.

The fact that the full-length dsbA(ss)-ExbD(44-63)-His6X construct was found in the cytoplasm was unexpected due to the co-translational localization preference of the dsbA(ss). In addition, it seems unlikely that the peptide would be inhibiting TonB function from the cytoplasm. The dsbA(ss)-ExbD(44-63) with or without the His tag was not captured with the mostly cytoplasmic residing TonB system membrane protein, ExbB. Considering that ExbB is the most abundant among the TonB system cytoplasmic membrane proteins, outnumbering ExbD 7:2 and TonB 7:1, and most of its protein is exposed to the cytoplasm (41), it seems unlikely that if this peptide were disrupting TonB system from the cytoplasm that it would not be doing so through contacting ExbB. Second, the ExbD and TonB small cytoplasmic portions can tolerate significant modifications. Half of the cytoplasmic portion of ExbD can be deleted and ToxR (residues 1-181) can be fused to the cytoplasmic portion of TonB without affecting TonB system activity (21, 31). Due to the modification tolerance in their small cytoplasmic domains of ExbD and TonB, it seems unlikely that either of these domains would serve as the peptide target. Taken together, it appears likely that the periplasmic portion of the peptide is inhibiting the TonB system.

### The ExbD disordered domain peptide complexes with ExbD and TonB are dependent on the presence of TonB and ExbD respectively

Three peptide-TonB complexes, two peptide-ExbD complexes, and one unknown ExbD complex were trapped with formaldehyde cross-linking from cells expressing dsbA(ss)-ExbD(44–63). At Stage I of TonB energization, ExbD and TonB do not interact. At Stage II, ExbD and TonB form a PMF-independent interaction and at Stage III, they form a PMF-dependent interaction (23, Kopp and Postle, manuscript submitted). To determine the stage of TonB energization at which the dsbA(ss)-ExbD(44-63) peptide interacts with ExbD and TonB, the ability for cells to form the peptide-complexes in absence of PMF, ExbD, or TonB was tested. The dsbA(ss)-ExbD(44-63) was expressed in the presence of the protonophore CCCP, in Δ*exbD* cells, and in Δ*tonB* cells (Fig. 6A and 6B), prior to formaldehyde treatment. The addition of CCCP, did not noticeably affect the peptide-ExbD complexes or the peptide-TonB complexes. This suggests that PMF is not required for the peptide to interact with either protein. In contrast, the absence of either TonB or ExbD prevented the peptide-ExbD and peptide-TonB complexes, respectively. The expression of dsbA(ss)-ExbD(44-63) captured significantly less ExbD-peptide_1_ and ExbD-peptide_2_ complexes when it was expressed in a *ΔtonB* strain compared when it was expressed in a WT strain (Fig. 6A). Similarly, dsbA(ss)-ExbD(44-63) was unable to capture TonB-peptide_1_B and TonB_1_C complexes when it was expressed in a *ΔexbD* strain (Fig. 6B). The deletion of either *tonB* or *exbD* did not significantly affect the peptide expression in either the whole cell and periplasm (Fig. 6C). Taken together, these data suggest that the dsbA(ss)-ExbD(44–63) is dependent on the presence of TonB and ExbD to interact with ExbD and TonB, respectively, but not PMF.

**Figure 6:**
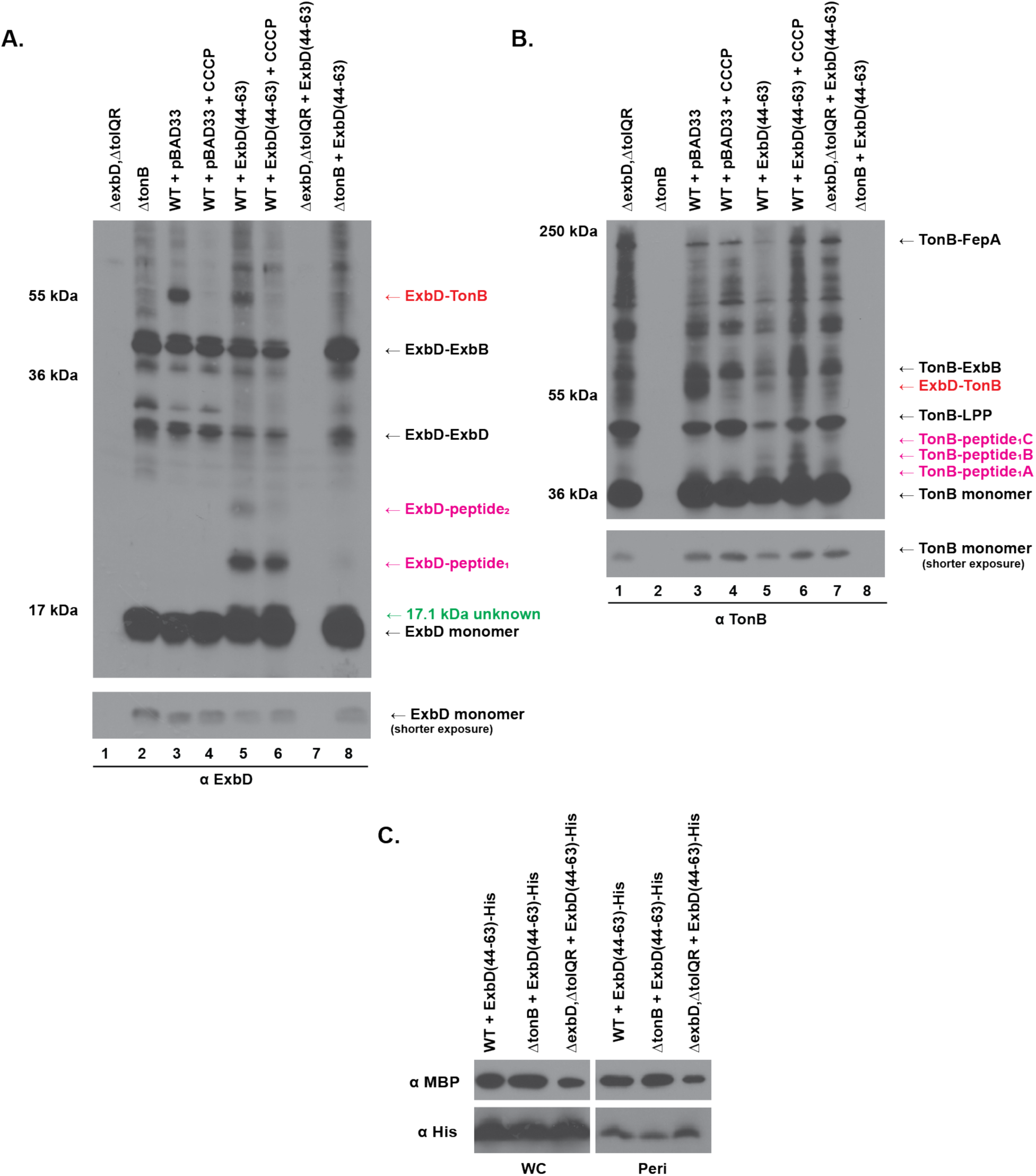
dsbA(ss)-ExbD(44-63) depends on the presence of ExbD and TonB to interact with TonB and ExbD respectively, but not PMF. Formaldehyde cross-linking of W3110 (WT) with a plasmid expressing of dsbA(ss)-ExbD(44-63) in the absence of PMF, and in the absence of the *exbD* gene, and in the absence of the *tonB* gene. Samples were probed with (A) anti-ExbD polyclonal antibodies or (B) anti-TonB monoclonal antibodies. Strains were grown to mid-exponential phase, at which point formaldehyde-cross-linking was performed as described in Materials and Methods. Equivalent numbers of cells were visualized on immunoblots of 13% SDS-polyacrylamide gels. (lane 1) ΔexbD, ΔtolQR refers to W3110 with the *exbD* gene and its homolog genes *tolQR* deleted (RA1045). (lane 2) ΔtonB refers to W3110 with the *tonB* gene deleted (KP1477). WT + pBAD33 refers to W3110 harboring the empty vector pBAD33 without (lane 3) and with (lane 4) the 60µM CCCP. WT + ExbD(44-63) is W3110 expressing the plasmid-encoded dsbA(ss)-ExbD(44-63) under the arabinose inducible promoter (pKP1847) with 0.2% (w/v) arabinose without (lane 4) and with (lane 5) the 60µM CCCP. (lane 6) ΔexbD, ΔtolQR + ExbD(44-63) refers to RA1045 expressing the plasmid-encoded dsbA(ss)-ExbD(44-63) under the arabinose inducible promoter (pKP1847) with 0.2% (w/v) arabinose. (lane 7) ΔtonB + ExbD(44-63) refers to KP1477 expressing the plasmid-encoded dsbA(ss)-ExbD(44-63) under the arabinose inducible promoter (pKP1847) with 0.2% (w/v) arabinose. (A, right) Positions of the previously characterized ExbD formaldehyde cross-linked complexes are shown: the PMF-dependent ExbD-TonB complex (red), the ExbD-ExbB complex, the ExbD homodimer (ExbD-ExbD), and ExbD monomer (25). (A, right labeled in purple) The suspected complexes of the dsbA(ss)-ExbD(44-63) peptide trapped with ExbD. (A, left) Mass markers are shown. (A, below) A shorter exposure of ExbD monomer corresponding to each sample. (B, right) Positions of the previously characterized TonB formaldehyde cross-linked complexes are shown TonB-FepA complex, the TonB-ExbB complex, the PMF-dependent TonB-ExbD complex, TonB-LPP (Braun’s lipoprotein) and TonB monomer (35, 25). (B, right labeled in purple) The suspected complexes of the dsbA(ss)-ExbD(44-63) peptide trapped with TonB. (left) Mass markers are shown. (B, below) A shorter exposure of TonB monomer corresponding to each sample. (C) Relative expression of plasmid-encoded dsbA(ss)-ExbD(44-63)-His6X expressed in W3110, KP1477, and RA1045 in whole cells (WC) and in the periplasm (Peri). Strains were grown to mid-exponential phase, at which point whole cells and soluble periplasmic samples were collected as described in the Materials and Methods. Equivalent numbers of cells were visualized on immunoblots of 16% acrylamide/ 6% bis-acrylamide SDS tricine gels. Protein expression was analyzed by probing with (Top panel) anti-maltose-binding protein (MBP) antibodies and (Bottom panel) anti-His monoclonal antibodies. The MBP, peptides, WC, periplasmic fractions were all from the same gel and the same immunoblot exposure for relative comparison. The plasmid identities are listed in Table S1.

### The first 5 residues within the peptide are important for the dsbA(ss)-ExbD(44–63) peptide to inhibit TonB system activity

To determine if the entire ExbD disordered domain was necessary for the dsbA(ss)-ExbD(44-63) to inhibit TonB system activity, truncations of the peptide were engineered: dsbA(ss)-ExbD(44-58)-His6X and dsbA(ss)-ExbD(49-63)-His6X. Although neither inhibited activity below 75% of WT, only the N-terminal truncation mutant, dsbA(ss)-ExbD(49-63), was as proteolytically stable as the parent peptide, dsbA(ss)-ExbD(44–63)-His6X (Fig. 7A, second from the bottom, and Fig. 7B, lane 4). The expression of the C-terminal truncation mutant, dsbA(ss)-ExbD(44–58)-His6X was undetectable (data not shown). Because dsbA(ss)-ExbD(49-63) was expressed similarly to the parent peptide but was significantly less inhibitory, it suggested that the first five residues of the dsbA(ss)-ExbD(44-63) peptide were important to inhibit TonB system activity.

**Figure 7:**
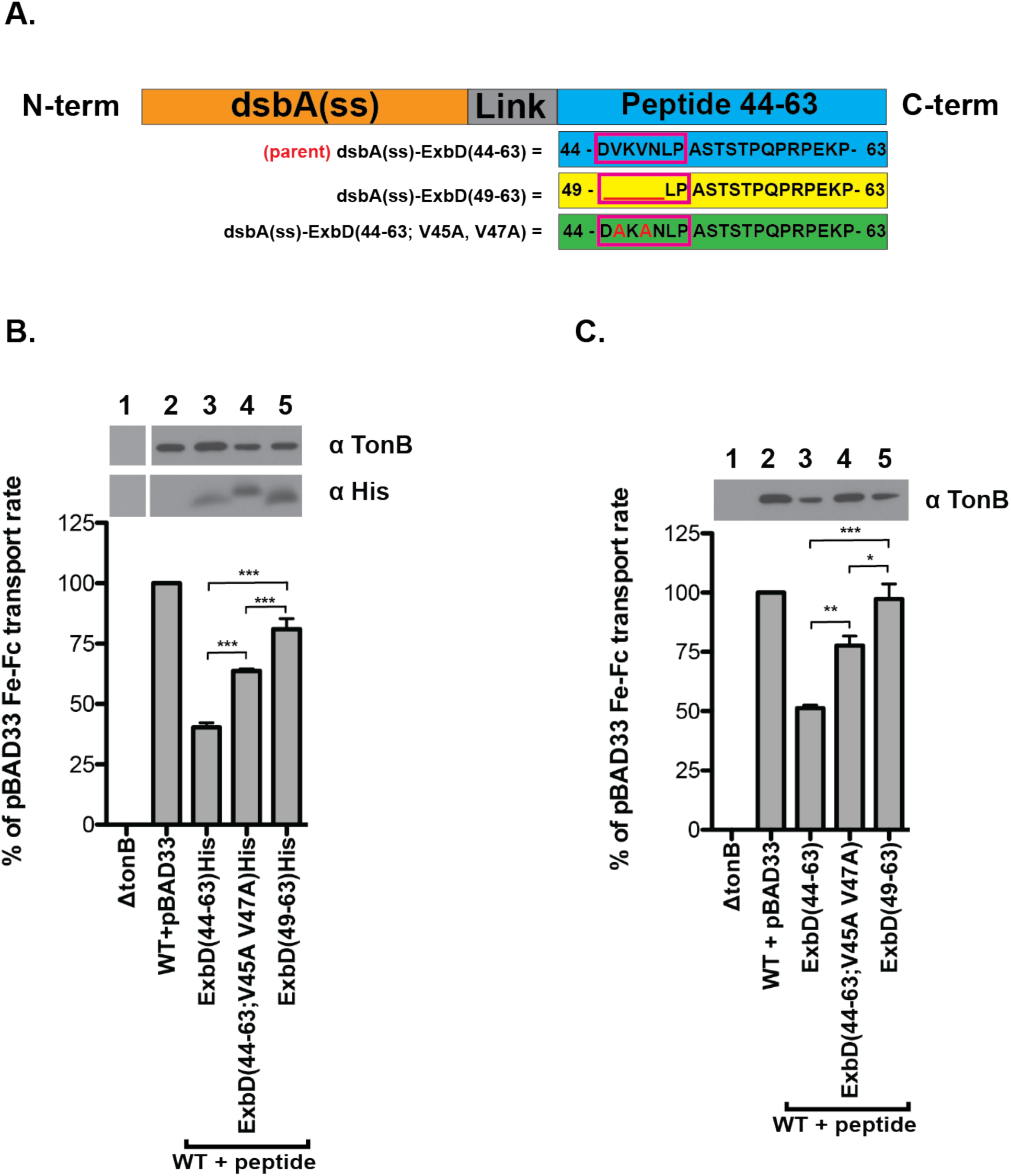
The first 5 residues of the ExbD(44-63) peptide including V45 and V47 are important for inhibiting TonB system activity. (A) Schematic representations of dsbA(ss)-ExbD(44-63) peptide derivatives were constructed. (A, top to bottom): The general representation of the parent dsbA(ss)-ExbD(44-63) construct. The ExbD amino acid sequence of the parent dsbA(ss)-ExbD(44-63) construct; the ExbD amino acid sequence with alanine substitutions for ExbD conserved valines within the dsbA(ss)-ExbD(44-63); a deletion of the first 5 residues within the dsbA(ss)-ExbD(44-63) construct. (B) Initial rate of ^55^Fe-ferrichrome transport of W3110 (WT) expressing derivatives of the inhibitory ExbD(44-63) peptide; ExbD(44-63; V45A, V47A) and ExbD(49-63) (B) with and (C) without the His6X-tag. Plasmids encoding ExbD (44-63), ExbD (44-63; V45A, V47A), and ExbD (49-63) were constructed with a cleavable, periplasmic localizing signal sequence, dsbA(ss) with and without a C-terminal-His6X-tag under the arabinose-inducible promoter. The dsbA signal sequence [dsbA(ss)] was fused to the N-terminus of ExbD(44-63) peptide for periplasmic localization, and four additional dsbA residues following the its peptidase cleavage site were added to minimize the potential cleavage interference from the exogenous peptide residues. The His6X tag was fused to the C-terminus of the ExbD(44-63) for peptide detection. Strains were grown to an A_550_ of 0.2 at which point 0.2% (w/v) of arabinose was added to induce intracellular expression of the peptides. Strains were grown until mid-exponential phase and the ^55^Fe-ferrichrome transport rate was measured as described in the Materials and Methods section. (above) The corresponding TCA precipitated protein samples were collected at the time of assay. Steady-state protein expression levels of TonB and the dsbA(ss)-ExbD(44-63)-His6X peptide (A only) were determined by immunoblot analysis with anti-TonB monoclonal antibodies and anti-His monoclonal antibodies, respectively. (left) The ^55^Fe-ferrichrome transport rate was recorded as the percent of the W3110 with the empty vector, pBAD33 (WT + pBAD33) transport rate. (lane 1) ΔtonB refers to W3110 with the *tonB* gene deleted (KP1477) and (lane 2) WT + pBAD33 refers to W3110 harboring the empty vector pBAD33. (A, lane 5) ExbD(44-63)His, ExbD(44-63; V45A, V47A)His, and ExbD(49-63)His refer to W3110 expressing the plasmid-encoded dsbA(ss)-ExbD(44-63)-His6X, dsbA(ss)-ExbD(44-63; V45A, V47A)-His6X, and dsbA(ss)-ExbD(49-63)-His6X, respectively, under the arabinose-inducible promoter with 0.2% arabinose (w/v). (B, lanes 3-5) ExbD(44-63), ExbD(44-63; V45A, V47A), and ExbD(49-63) refer to W3110 expressing the plasmid-encoded dsbA(ss)-ExbD(44-63), dsbA(ss)-ExbD(44-63; V45A, V47A), and dsbA(ss)-ExbD(49-63), respectively, under the arabinose-inducible promoter with 0.2% arabinose (w/v). AVONA analysis followed by Tukey’s multiple comparison test was used to measure significance from 3 independent experiments performed in triplicate. The 3 asterisks indicate a p-value of < 0.001. The plasmid identities are listed in Table S1.

Previously, we have identified a conserved ΨXΨXLP motif within the ExbD disordered domain that was essential for TonB system activity (Kopp and Postle, manuscript submitted). We also had demonstrated that alanine substitutions for the conserved hydrophobic residues (valines 45 and 47 in *Escherichia coli*) within the motif prevented distal ExbD interactions with TonB (Kopp and Postle, manuscript submitted). Within the context of the dsbA(ss)-ExbD(44-63) peptide the first five residues, which consists of ExbD conserved valines, V45 and V47, were found to be important for the peptide to inhibit TonB system activity. To determine whether the valines were important for the dsbA(ss)-ExbD(44-63) peptide to inhibit TonB activity, we constructed a version of the peptide with alanine substitutions for peptide residues V45 and V47 [dsbA(ss)-ExbD(44-63;V45A, V47A) (Fig. 7A, bottom).

Cells expressing dsbA(ss)-ExbD(44-63; V45A, V47A) + or - the His-tag inhibited Fe-transport significantly less than those expressing dsbA(ss)-ExbD(44-63) + or - the His-tag (Fig. 7B and 7C, lanes 4 and 3) However, cells expressing dsbA(ss)-ExbD(49-63) + or - the His-tag were significantly less inhibitory than dsbA(ss)-ExbD(44-63; V45A, V47A) + or - the His tag (Fig. 7B and 7C, lanes 5 and 4). This suggests that although valines 45 and 47 contribute to the dsbA(ss)-ExbD(44–63) inhibiting TonB activity, they are not the only important residues within the first five. Albeit the His-tagged versions of each peptide were ∼10% more inhibitory than those without, the relative TonB inhibitory difference was similar between the peptides + or - the His-tags.

### The ExbD(49-63) and ExbD(44-63; V45A, V47A) peptide variants did not inhibit the PMF-dependent ExbD-TonB interaction and interact differently with ExbD and TonB than the inhibitory peptide, dsbA(ss)-ExbD (44-63)

The effect on TonB system and ExbD-peptide and TonB-peptide complexes from cells expressing the peptide variants, dsbA(ss)-ExbD(44-63; V45A, V47A) and dsbA(ss)-ExbD(49-63) were compared to that of the dsbA(ss)-ExbD(44-63) peptide. The expression of either dsbA(ss)-ExbD(49-63) or dsbA(ss)-ExbD(44-63; V45A, V47A) did not significantly inhibit ExbD or TonB complexes, including that of the PMF-dependent ExbD-TonB interaction (Fig. 8A and 8B). In addition, the peptide variants, dsbA(ss)-ExbD(49-63) or dsbA(ss)-ExbD(44-63; V45A, V47A) still captured in ExbD-peptide_1_ and ExbD-peptide_2_ formaldehyde cross-linked complexes, although not as significantly as dsbA(ss)-ExbD(44-63). In respect to TonB-peptide complexes, dsbA(ss)-ExbD(44-63; V45A, V47A) captured noticeably lessTonB-peptide1B and TonB-peptide1C complexes than dsbA(ss)-ExbD(44-63) whereas dsbA(ss)-ExbD(49-63) did not trap any detectable TonB-peptide_1_B and TonB-peptide_1_C complexes. These data demonstrate that the peptide variants interact differently with ExbD or TonB compared to the parent inhibitory peptide, dsbA(ss)-ExbD(44-63).

**Figure 8:**
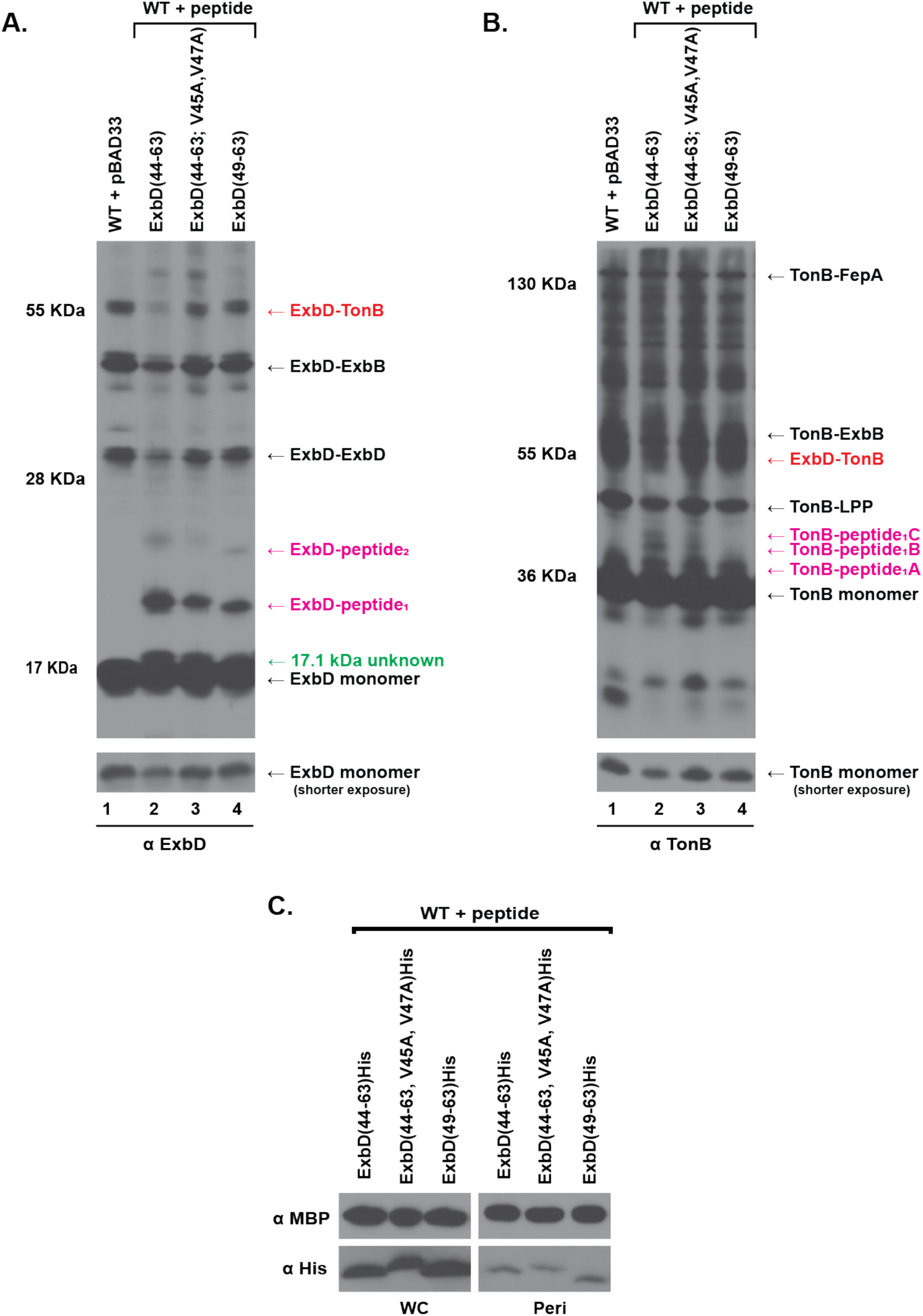
dsbA(ss)-ExbD(44-63; V45A, V47A) and dsbA(ss)-ExbD(49-63) do not inhibit the ExbD-TonB complex and reduced TonB interactions. Formaldehyde cross-linking of W3110 (WT) expressing the dsbA(ss)-ExbD(44-63), dsbA(ss)-ExbD(44-63; V45A, V47A) or dsbA(ss)-ExbD(49-63) probed with (A) anti-ExbD polyclonal antibodies or (B) anti-TonB monoclonal antibodies. Strains were grown to mid-exponential phase, at which point formaldehyde-cross-linking was performed as described in Materials and Methods. Equivalent numbers of cells were visualized on immunoblots of 13% SDS-polyacrylamide gels. (A, top) (lane 1) WT + pBAD33 refers to W3110 harboring the empty vector pBAD33 ExbD(44-63), (lane 2) ExbD(44-63), (lane 3) ExbD(44-63; V45A, V47A), and (lane 4) ExbD(49-63) refer to W3110 expressing the plasmid-encoded dsbA(ss)-ExbD(44-63), dsbA(ss)-ExbD(44-63; V45A, V47A), and dsbA(ss)-ExbD(49-63). (A, right) Positions of the previously characterized ExbD formaldehyde cross-linked complexes are shown: the PMF-dependent ExbD-TonB complex (red), the ExbD-ExbB complex, the ExbD homodimer (ExbD-ExbD), and ExbD monomer (2). (right, labeled in purple) The suspected complexes of the dsbA(ss)-ExbD(44-63) peptide trapped with ExbD. (right in green) identifies the 17.1 kDa unknown ExbD complex (A, right) Mass markers are shown. (A, bottom) A shorter exposure of ExbD monomer corresponding to each sample. (B, top) (lane 1) WT + pBAD33 refers to W3110 harboring the empty vector pBAD33, (lane 2) ExbD(44-63), (lane 3) ExbD(44-63; V45A, V47A), and (lane 4) ExbD(49-63) refer to W3110 expressing the plasmid-encoded dsbA(ss)-ExbD(44-63)-His6X, dsbA(ss)-ExbD(44-63; V45A, V47A)-His6X, and dsbA(ss)-ExbD(49-63)-His6X. (B, right) Positions of the previously characterized TonB formaldehyde cross-linked complexes are shown TonB-FepA complex, the TonB-ExbB complex, the PMF-dependent TonB-ExbD complex (red), TonB-LPP (Braun’s lipoprotein) and TonB monomer (3, 2). (B, right, labeled in purple) The suspected complexes of the peptide trapped with TonB: TonB-peptide1A, TonB-peptide1B, and TonB-peptide1C. (B, left) Mass markers are shown. (B, bottom) A shorter exposure of TonB monomer corresponding to each sample. (C) Relative expression of plasmid-encoded dsbA(ss)-ExbD(44-63)-His6X, dsbA(ss)-ExbD(44-63; V45A, V47A)-His6X, and dsbA(ss)-ExbD(49-63)-His6X in in whole cells (WC) and in the periplasm (Peri). Strains were grown to mid-exponential phase, at which point whole cells and soluble periplasmic samples were collected as described in the Materials and Methods. Equivalent numbers of cells were visualized on immunoblots of 16% acrylamide/ 6% bis-acrylamide SDS tricine gels. Protein expression was analyzed by probing with (Top panel) anti-maltose-binding protein (MBP) antibodies and (Bottom panel) anti-His monoclonal antibodies. The MBP, peptides, WC, periplasmic fractions were all from the same gel and the same immunoblot exposure for relative comparison. The plasmid identities are listed in Table S1.

Interestingly, the His-tagged versions of the peptides were captured differently with ExbD and TonB than the those without (Fig. 9). The dsbA(ss)-ExbD(44-63)-His6X peptide captured less of the ExbD-pepitde1 complex than either dsbA(ss)-ExbD(44-63; V45A, V47A)-His 6X or dsbA(ss)-ExbD(49-63)-His6X. However, it captured more of the 17.1 kDa unknown ExbD complex than dsbA(ss)-ExbD(49-63)-His6X, but similar to that of dsbA(ss)-ExbD(44-63; V45A, V47A)-His6X. The peptides also trapped with TonB differently. The dsbA(ss)-ExbD(44-63)-His6X peptide trapped all three of TonB-peptide complexes less than the version without the His-tag. But, also dsbA(ss)-ExbD(44-63)-His6X peptide trapped all three less than dsbA(ss)-ExbD(44-63; V45A, V47A)-His6X. In comparison to dsbA(ss)-ExbD(49-63)-His6X, dsbA(ss)-ExbD(44-63)-His6X peptide trapped less TonB-peptide_1_B and TonB-peptide_1_C complexes but similar amounts of TonB-peptide_1_A.

**Figure 9:**
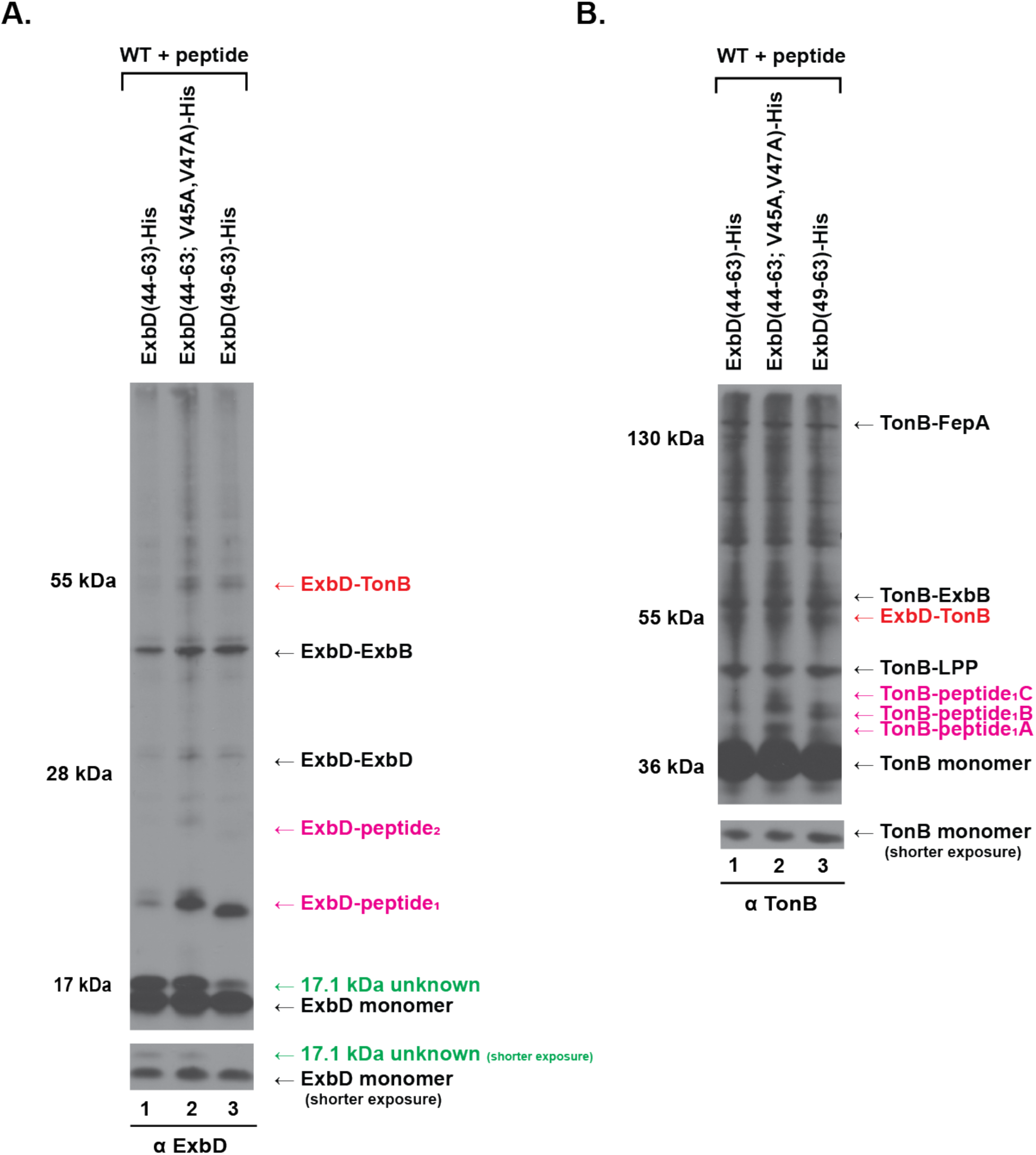
The His-tag alter formaldehyde cross-linking of dsbA(ss)-ExbD(44-63) peptides to ExbD and TonB. Formaldehyde cross-linking of W3110 (WT) expressing the dsbA(ss)-ExbD(44-63)-His6X, dsbA(ss)-ExbD(44-63; V45A, V47A)-His6X or dsbA(ss)-ExbD(49-63)-His6X probed with (A) anti-ExbD polyclonal antibodies or (B) anti-TonB monoclonal antibodies. Strains were grown to mid-exponential phase, at which point formaldehyde-cross-linking was performed as described in Materials and Methods. Equivalent numbers of cells were visualized on immunoblots of 13% SDS-polyacrylamide gels. (A, top) (lane 1) ExbD(44-63)-His, (lane 2) ExbD(44-63; V45A, V47A)-His, and (lane 3) ExbD(49-63)-His refer to W3110 expressing the plasmid-encoded dsbA(ss)-ExbD(44-63)-His6X, dsbA(ss)-ExbD(44-63; V45A, V47A)-His6X, and dsbA(ss)-ExbD(49-63)-His6X. (A, right) Positions of the previously characterized ExbD formaldehyde cross-linked complexes are shown: the PMF-dependent ExbD-TonB complex (red), the ExbD-ExbB complex, the ExbD homodimer (ExbD-ExbD), and ExbD monomer (2). (right, labeled in purple) The suspected complexes of the dsbA(ss)-ExbD(44-63) peptide trapped with ExbD. (right in green) identifies the 17.1 kDa unknown ExbD complex (A, right) Mass markers are shown. (A, bottom) A shorter exposure of ExbD monomer and the 17.1 kDa unknown ExbD complex corresponding to each sample. (B, top) (lane 1) ExbD(44-63)-His, (lane 2) ExbD(44-63; V45A, V47A)-His, and (lane 3) ExbD(49-63)-His refer to W3110 expressing the plasmid-encoded dsbA(ss)-ExbD(44-63)-His6X, dsbA(ss)-ExbD(44-63; V45A, V47A)-His6X, and dsbA(ss)-ExbD(49-63)-His6X. (B, right) Positions of the previously characterized TonB formaldehyde cross-linked complexes are shown TonB-FepA complex, the TonB-ExbB complex, the PMF-dependent TonB-ExbD complex (red), TonB-LPP (Braun’s lipoprotein) and TonB monomer (3, 2). (B, right, labeled in purple) The suspected complexes of the peptide trapped with TonB: TonB-peptide1A, TonB-peptide1B, and TonB-peptide1C. (B, left) Mass markers are shown. (B, bottom) A shorter exposure of TonB monomer corresponding to each sample.

Because the His-tags on the peptides did not alter their relative effect on TonB-dependent ^55^Fe-transport (Fig. 7B and 7C), they likely do not affect how each peptide affects TonB system function. However, differences in formaldehyde cross-linked ExbD and TonB complexes trapped among the peptides + or - the His-tags likely demonstrate the variation in which peptides interact with ExbD and TonB differently. No peptide-ExbB complex was identified in cells expressing any version of peptide, + or - the His-tags (Fig. S6).

### The ExbD disordered domain peptide affects PMF integrity

The TonB system requires PMF to function. To test whether the peptides affect the integrity of PMF, the ability for cells to expel ethidium bromide (EtBr) via the TolC efflux pump was measured. The accumulation of EtBr in the cells is inversely proportional to the PMF. Although none of the peptides affected PMF to the degree at which would inhibit TonB system activity (Fig. 10A and 10B), there was a correlation between proton leakage and TonB system inhibition caused from expressing the peptides (Fig. 10C and 10D). The expression of dsbA(ss)-ExbD(44–63) with and without the His-tag caused cells to leak protons significantly more than dsbA(ss)-ExbD(44-63; V45A, V47A) with and without the His-tag or dsbA(ss)-ExbD(49–63) with and without the His-tag. However, the degree at to which dsbA(ss)-ExbD(44–63) leaked protons was not enough to contribute to TonB system activity (Fig. 10B). WT cells treated with 15μM CCCP accumulated EtBr at approximately double the rate of than cells expressing dsbA(ss)-ExbD(44–63)-His6X, did not have any explain any of the loss in TonB system activity (Fig. 10B). Therefore, the leaking of protons cellularly cannot explain the peptides ability to inhibit the TonB system. However, the correlation between proton leakage caused by the expression of the peptides is difficult to ignore. Similar observations were made previously with mutations within ExbD the conserved ΨXΨXLP motif. But it was unclear of the connection between the PMF and the ExbD ΨXΨXLP motif. Perhaps, the dsbA(ss)-ExbD(44–63) peptide may be interacting with ExbD or TonB in way that causes a localized proton leakage through the TonB system. However, the knowledge of how or why this would occur is not clear.

**Figure 10:**
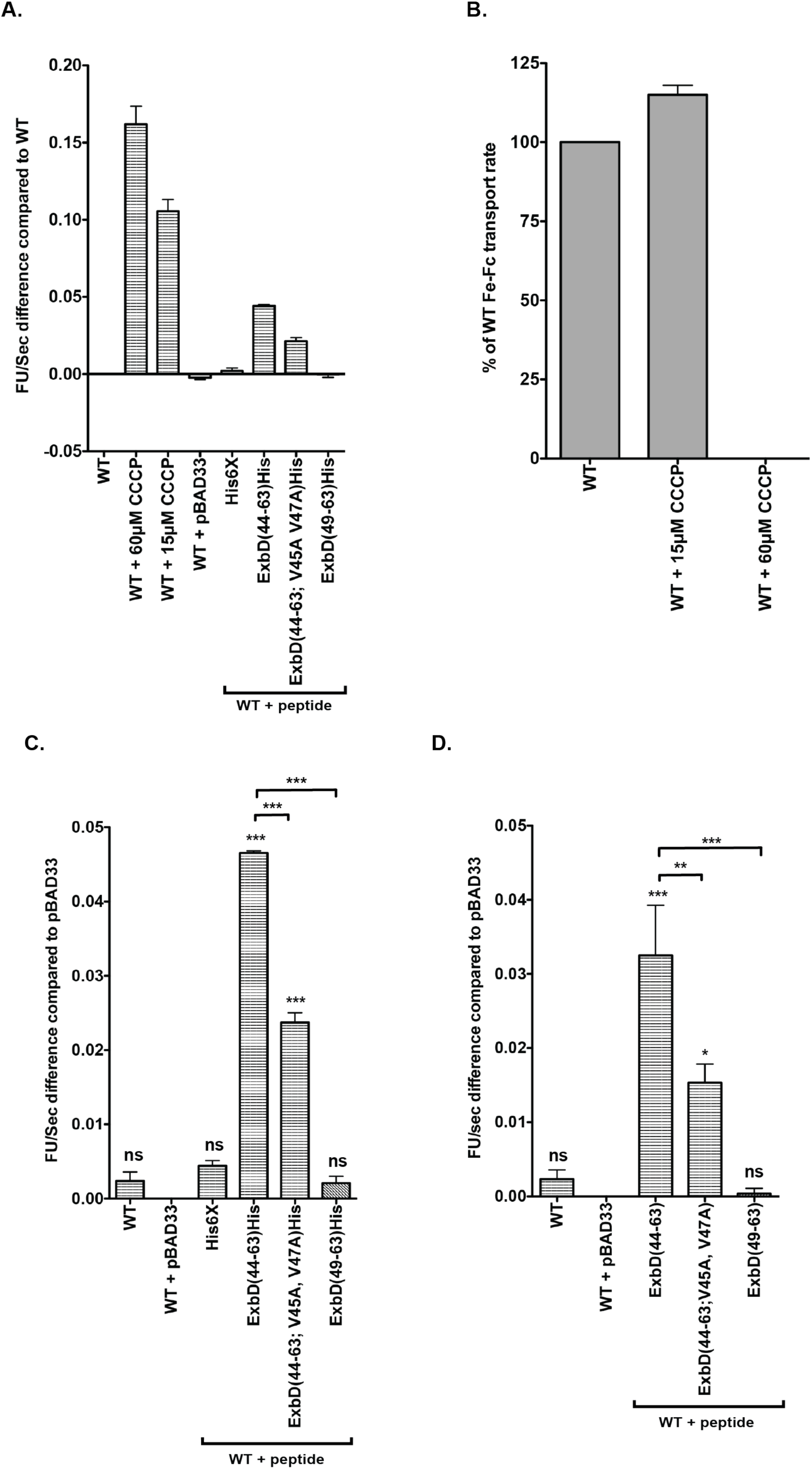
dsbA(ss)-ExbD(44-63) significantly leaks protons but not enough systematically to affect TonB system activity. (A) EtBr accumulation assay for W3110 in the presence of the protonophore, CCCP and expressing plasmid-encoded peptides with His6X tags. Strains were grown to an A_550_ of 0.2 at which point 0.2% (w/v) of arabinose was added to induce intracellular expression of the peptides. The strains were grown until mid-exponential phase and the rate of ethidium bromide accumulation was measured as described in the Materials and Methods section. (left) The ethidium bromide accumulation rate, measured as fluorescent units / second (FU/sec), was recorded as the rate difference to that of the W3110 with the empty vector, pBAD33 (WT + pBAD33) transport rate. (bottom) WT refers to W3110, and WT + 60 µM CCCP and WT +15 µM CCCP refer to W3110 treated with the respected concentrations of the protonophore, CCCP, prior to measuring the ethidium accumulation. WT + pBAD33 refers to W3110 harboring the empty vector pBAD33. His6X refers to W3110 expressing a plasmid-encoded dsbA signal sequence plus the four amino-acid linker sequence fused to C-terminal-His6X under the arabinose-inducible promoter (pKP1977) with 0.2% (w/v) arabinose. ExbD(44-63)His, ExbD(44-63; V45A, V47A)His, and ExbD(49-63)His refer to W3110 expressing the plasmid-encoded dsbA(ss)-ExbD(44-63)-His6X, dsbA(ss)-ExbD(44-63; V45A, V47A)-His6X, and dsbA(ss)-ExbD(49-63)-His6X, respectively, under the arabinose-inducible promoter with 0.2% arabinose (w/v). The plasmid identities are listed in Table S1. (B) Initial rate of ^55^Fe-ferrichrome transport of W3110 (WT) in the presence of 15 µM and 60 µM CCCP. Strains were grown until mid-exponential phase and the ^55^Fe-ferrichrome transport rate was measured as described in the Materials and Methods section. (left) The ^55^Fe-ferrichrome transport rate was recorded as the percent of the W3110 (WT) transport rate. WT + 60 µM CCCP and WT +15 µM CCCP refer to W3110 treated with the respected concentrations of the protonophore, CCCP, at the time of the assay (C) EtBr accumulation assay for W3110 expressing plasmid-encoded peptides with His6X tags. Samples and data are the same from (A) without the CCCP treated samples. (D) EtBr accumulation assay for W3110 expressing plasmid-encoded peptides without His6X tags. Experiment and procedure were performed identically as (A) and (C). AVONA analysis followed by followed by Tukey’s multiple comparison test was used to measure significance from three independent experiments. The *** indicates a p-value of < 0.001, ** indicates a p-value of < 0.01, * indicates a p-value of < 0.05, and ns is not significant. The symbols immediately above the error bars are as they compare to WT + pBAD33. The symbols above the brackets refer to the comparison between the two specific samples indicated by each bracket. The plasmid identities are listed in Table S1.

## DISCUSSION

The TonB system is a virulence factor for many Gram-negative pathogens, mostly for iron acquisition (4–6, 42). Because the TonB system is unique to Gram-negative bacteria with no homologous system host, it is an appealing target for a novel antibiotic. However, no antibiotic to date exists against this system. Here, we have demonstrated that an ExbD(44-63) peptide can inhibit the ExbD complexes including the ExbD-TonB PMF-dependent interaction. Mutations within the peptide conserved ExbD ΨXΨXLP motif had rendered the peptide significantly less inhibitory. The importance of the ΨXΨXLP motif within the inhibiting peptide may serve as a foundation to develop a novel antibiotic against the system.

### Making sense of the formaldehyde cross-linked peptide complexes

The dsbA(ss)-ExbD(44-63) peptide expression caused the formation of three unknown TonB formaldehyde cross-linked complexes and three unknown ExbD formaldehyde cross-linked complexes. The TonB unknown complexes were identified as three distinct TonB with one peptide complexes (TonB-peptide_1_A, TonB-peptide_1_B, and TonB-peptide_1_C) whereas the ExbD unknown complexes were identified as an ExbD with one peptide (ExbD-peptide_1_), an ExbD-peptide with two peptides (ExbD-peptide_2_), and an unknown ExbD 17.1 kDa complex that did not contain a peptide. All the TonB-peptide complexes were not present without ExbD. Similarly, ExbD-peptide_2_ complex was not present and ExbD-peptide_1_ complex was noticeably reduced in the absence of TonB. However, none of the complexes were affected in absence of PMF. Because ExbD and TonB are required to interact PMF-independently at Stage II in the TonB energization cycle and PMF-dependently at Stage III (23), it suggested that the dsbA(ss)-ExbD(44-63) interacted with ExbD and TonB after Stage II, but before Stage III. In contrast, the ExbD 17.1 kDa unknown complex was not affected in the absence of TonB which suggested that this complex formed prior to Stage II.

Previously, disulfide cross-linking experiments showed that TonB formed three distinct complexes of TonB homodimers, which are referred to as the TonB triplet-homodimers (43). The formation of the triplet TonB homodimers required a functional TonB TMD and ExbBD (43). Two of three TonB homodimers required ExbD (15). This suggested that at least two of three TonB homodimers occur after Stage II. Considering that the dsbA(ss)-ExbD(44-63) peptide interacted with TonB to form three separate complexes during Stage II, it may be possible that these complexes are related to the multiple conformations of TonB homodimeric complexes during Stage II. Because disordered domains can assume multiple protein conformations and possibly induce conformational changes in its binding partners (44, 45), the ExbD disordered domain may mediate TonB homodimeric transitions. Further speculation would suggest that the ExbD conserved ΨXΨXLP motif specifically would dictate these transitions. In support of this hypothesis, is that the expression of the dsbA(ss)-ExbD(44-63) derivatives, dsbA(ss)-ExbD(49-63) and ExbD(44-63; V45A, V47A) altered the TonB-peptide formaldehyde cross-linked complexes, which suggests that the motif dictates how the peptides interact with TonB.

Two dsbA(ss)-ExbD(44-63) peptide-ExbD complexes were also identified from formaldehyde cross-linking: ExbD-peptide_1_, which consisted of ExbD and a single peptide, and ExbD-peptide_2_, which consisted of ExbD and two peptides. Both ExbD-peptide complexes were dependent on TonB but not PMF. This may be evidence of ExbD homodimeric interactions that occur after Stage II. It has been well documented that ExbD homodimers form regardless if TonB is present (25, 29, 30, 46). Previously, *in vivo* photo-cross-linking captured ExbD homodimers throughout the ExbD disordered domain. However, the technique was unable to distinguish from one homodimeric interaction from another. Because that two peptides, albeit weakly, were captured in a complex with ExbD, this may suggest that either that the peptides interact with ExbD in two separate locations or that the two peptides interact together at the same location on ExbD. However, the latter would suggest of an ExbD trimeric interaction and there is no current *in vivo* evidence supporting the existence of such a complex (2, Kopp and Postle, manuscript submitted). Furthermore, if the ExbD-peptide_2_ complex formed from non-specific aggregation, then one would probably expect to see higher-order ExbD aggregates with the peptides rather than only a complex with two-peptides and ExbD. Therefore, the idea of the two peptides interacting in at least two different locations on ExbD appears to be the more likely one of the two scenarios.

So where would the two locations be in a scenario where the ExbD disordered domain interacted in two locations? ExbD di-sulfide cross-linked homodimers were captured through identical residues through-out the disordered region (Fisher et al. unpublished data). Yet, *in vivo* photo-cross linking from the ExbD disordered domain could not rule out the possibility of a beta-strand swap ExbD dimeric interaction that would have been predicted from a TolR dimeric crystal structure that had been published (47). The ExbD homodimeric photo-cross-linked complexes captured from residues within the ExbD disordered domain varied in which the most captured homodimers correlated mostly, although not identically, with the residues that would be predicted from the TolR dimeric crystal structure. However, these same residues were also captured via disulfide cross-linking through identical residues instead a beta-strand swap. Perhaps, the higher amount of the ExbD homodimer photo-cross-linked complexes may correlate with the different homodimeric interactions. However, more direct evidence will be needed to verify the existence of beta-swap conformation of ExbD *in vivo* to support these speculations.

The two peptide-ExbD complexes appear also to be dependent following Stage II, but prior to Stage III. Interestingly though, no ExbD homodimeric interaction has been conclusively shown to exist that is dependent on TonB (25, 29, 30, 46) which may suggest that these ExbD-peptide interactions are indicative of novel ExbD homodimeric interactions. Although the identical residues within the ExbD disordered domain that formed di-sulfide bridges were not dependent on TonB (Fisher et al. unpublished data), the presence of a potential beta-strand swap ExbD homodimer and whether it is dependent on TonB *in vivo* has not been confirmed. It is possible that this homodimeric conformation may be dependent of TonB *in vivo*. Moreover, alanine substitutions within the conserved ΨXΨXLP motif within the ExbD disordered domain, altered the formaldehyde crosslinking of the ExbD dimer to form three complexes (Kopp and Postle, manuscript submitted). Considering that the motif is only required to transition from Stage II to Stage III, this may support a mechanism of the motif that is involved in ExbD homodimeric interactions at or after Stage II. However, it is unclear when the Stage II ExbD dimeric interactions would occur in relation to the ExbD-TonB PMF-independent interactions, but the evidence presented in this work suggests potential substages between Stage II and Stage III.

### The potential identification of the ExbD unknown 17.1 kDa complex

Although no peptide was detected in the unknown ExbD 17.1 kDa complex, it was dependent on peptide expression. The complex is only ∼1 kDa more than ExbD monomer, which indicated that this may be not be a complex with another protein, but it may be an ExbD conformational change. In this scenario, the peptide would interact with ExbD causing a conformational change, but a complex would not be captured because the formaldehyde cross-linkable residues do not align. To support this argument, ExbD pBpa substitutions and alanine substitutions for residues within the motif cause a monomer doublet, with the top band in the doublet being at a similar molecular weight distance as the 17.1 unknown complex is to the ExbD monomer. Therefore, the 17.1 kDa may be a conformational change in ExbD that can be captured with formaldehyde and occurs prior to Stage II.

### ExbD(44–63) may not be the only ExbD inhibitory peptide

Although dsbA(ss)-ExbD(44–63) may serve as a template to potentially design inhibitors against the TonB system, there may be other candidates to explore. One specifically is the ExbD(112–131) peptide. Although the steady-state levels were relatively low, it still prevented TonB system activity to about 70% of WT. An amphipathic helix (ExbD residues 112-125) is present within this region for which its function remains unclear in ExbD is unclear. The ExbD homolog, TolR, amphipathic helix has been suggested to serve as “a plug-like” function that controls proton flow, but the evidence is not conclusive (48). Unfortunately, the dsbA(ss)-ExbD(112–131) steady-state levels were not adequate to detect significant changes in any ExbD or TonB complex (data not shown). However, it would be interesting if proteolytically stable alterations could be made to the dsbA(ss)-ExbD(112–131) peptide to possibility identify the target(s) of this peptide.

### The correlation of the disordered domain peptide and proton leakage

The ExbD homolog, MotB has a periplasmic “plug” domain which is hypothesized to control proton flow. However, no equivalent domain exists in ExbD or TolR. Here, the dsbA(ss)-ExbD(44-63) with or without the His-tag causes significant proton leakage, but the systematic leakage was not enough to inhibit TonB activity. The idea that the leakage was merely a correlation and was not related to TonB inhibition seems unlikely considering that the proton leakage directly correlated with activity loss. Moreover, the idea that the peptides caused proton leakage locally via the TonB system and/or possibly the Tol system also does not appear to be likely because preliminary evidence suggests that the proton leakage of dsbA(ss)-ExbD(44-63)-His6X, still was significant in the absence of the TonB and Tol System (data not shown). This would suggest, although maybe not entirely, most of the leakage is independent of the systems. Instead, it may indicate that the intrinsic conformation of the ExbD disordered domain may initiate protons through the membranes. The first five residues of the peptide are required for proton leakage of the peptide, which would place these residues near the cytoplasmic membrane-periplasmic interface. Interestingly, dsbA(ss)-ExbD(44-141)-His did not leak protons which may suggest that the C-terminal domain may regulate the possible function of the disordered domain controlling protons. However, further experiments would be needed to clarify the connection between proton leakage and the ExbD disordered domain.

### Why the ExbD and TonB soluble periplasmic domains formaldehyde cross-link to TonB and ExbD in the absence of PMF

The periplasmic domains of ExbD and TonB have each shown to inhibit TonB-dependent activity and prevent the ExbD-TonB PMF-dependent interaction. The PMF-dependent ExbD-TonB interaction can be captured using *in vivo* formaldehyde cross-linking (25) but, in the absence of PMF, ExbB, or their TMDs, ExbD and TonB do not form a formaldehyde cross-linked complex (23, 25, 49–52). However, here we showed that the periplasmic domains of TonB or ExbD that were fused to cleavable signal sequences formed formaldehyde cross-linked complexes with ExbD and TonB, respectively. This may suggest that PMF, ExbB, and TMDs may articulately control the ExbD conformations rather than be required for ExbD to assume a conformation that formaldehyde cross-links with TonB. However, because the periplasmic domains alone are not sufficient to support TonB system activity, there are other factors contribute to energizing TonB directly. Indeed, no known example has been shown where the ExbD-TonB PMF-dependent interaction is present and the TonB system is lost. But it may be possible that a subsequent sub-stage of TonB energization involving the ExbD and TonB TMDs following the PMF-dependent ExbD-TonB interaction but prior for TonB to transduce the energy to the transporters. Alternatively, because the ExbD and TonB periplasmic domains require higher expression than chromosomal ExbD and TonB to be captured with formaldehyde, it may be possible that the PMF-independent ExbD-TonB interaction can be captured with formaldehyde, but the crosslinking is too weak to be detected at chromosomal levels. More information will be needed to discern these two possibilities.

### Summary

This work demonstrated that the PMF-dependent ExbD-TonB interaction could be inhibited by either the TonB periplasmic domain or the ExbD periplasmic domain. Furthermore, the ExbD disordered domain was captured in multiple complexes with TonB and ExbD; many of which appeared to have formed during Stage II to Stage III of TonB energization. The conserved motif is a key feature within the dsbA(ss)-ExbD(44-63) peptide in which mutations or partial deletions within the motif relieved its inhibitory potency. The data further suggested that the motif-deficient versions of the dsbA(ss)-ExbD(44-63) peptide interact differently with TonB and ExbD. Considering the importance of the ExbD conserved motif and its ability to disrupt protein-protein interactions within the TonB system, it may serve as a potential novel antibiotic target or a potential design for an inhibitor against the TonB system.

## MATERIALS AND METHODS

### Strains and plasmids

Strains and plasmids used in this study are listed in Table S1. The plasmids that were used were those with a pBAD33 backbone. pKP1715 and pKP1714 were constructed via cloning by BamHI restriction digest and ligation of the ompT signal sequence with TonB(33-239) and the ompT signal sequence with ExbD(44-141) from pKP1695 and pKP1694 into pBAD33 vectors. pKP1695 and pKP1694 were constructed via cloning by BamHI restriction digest and ligation of TonB(33-239) and ExbD(44-141) from pKP325 and from pKP999 into pET12a vectors, which have the ompT leader signal sequence under the T7 promoter. The ompT linker sequence was a byproduct of restriction digest and ligation with BamHI that added four addition codons downstream of its peptidase cleavage site (Fig. S1).

The plasmid, pKP1832, was constructed from using a two-step PCR process to swap the ompT signal sequence and linker sequence in pKP1714 for the dsbA signal sequence [dsbA(ss)] plus four additional dsbA amino-acids downstream of its signal peptidase cleavage site. First, the dsbA(ss) was amplified from the ASKA dsbA gene, in which the 3’ prime halves of the forward and reverse primers were complementary to the dsbA(ss). The 5’ prime haves of the primers were designed in a way that the amplified dsbA(ss) product from the first round of PCR served as “primers” a second round of PCR to replace the ompT(ss) with the dsbA(ss) in pKP1714. All of dsbA(ss) plasmids were constructed by quick-change PCR mutagenesis using 25-cycle extralong PCR with 5’-phosphorylated primers. pKP1838, pKP1839, pKP1977, and pKP2013 were derived from pKP1832. pKP1847 and pKP1946 were derived from pKP1838 whereas pKP1855 and pKP1856 were derived from pKP1839. pKP1960 and pKP2089 were derived from pKP1847. pKP1961, pKP1980, and pKP2006 were derived from pKP1946, pKP1855, and pKP1856, respectively. pKP1981, pKP1982, and pKP1983 were all derived from pKP1961. pKP2142 and pKP2069 were derived from pKP1960. pKP2082 was derived from pKP2069. All mutants were verified by DNA sequencing at the Penn State Genomics Core Facility, University Park, PA.

### Initial rates of **^55^**Fe-ferrichrome transport

Cultures were grown in Luria-Bertani at 37°C with aeration to saturation. The saturated cultures were sub-cultured in were 1:100 in M9 minimal media supplemented with 0.4% glycerol, 0.2% Casamino Acids, 40 µg/ml tryptophan, 4 µg/ml vitamin B1, 1 mM MgSO4, 0.5 mM CaCl2, and 37 µM FeCl3 and grown until mid-exponential phase. Strains harboring the pBAD33 backbone plasmids were treated with 0.2% arabinose at A_550_ of ∼0.2 to induce protein expression. For detecting the relative peptide expressions for cells harboring pKP2013 in the ^55^Fe-ferrichrome assay of different concentrations of 0.0025%, 0.005%, 0.01%, 0.02%, 0.05%, and 0.2% arabinose were used. The ^55^Fe-ferrichrome assay was performed in triplicate from at least two independent experiments as previously described (53). To determine steady state levels of TonB and the His-tagged peptides, samples were precipitated with 10% TCA at the start of the assay and equivalent numbers of cells processed for immunoblot analysis on either 13% SDS-polyacrylamide glycine gels or 16% acrylamide with 6% bis-acrylamide tricine gels (56). TonB and the His-tagged peptides were detected by immunoblotting with anti-TonB 4F1 monoclonal antibodies (57), and anti-His monoclonal antibodies [EMD Millipore, prod # 70796].

### Iron-dependent growth curve

Cultures were grown in Luria-Bertani at 37°C with aeration to saturation. Each culture was sub-cultured in 1:1000 in M9 minimal media supplemented with 0.4% glycerol, 0.2% Casamino Acids, 40 µg/ml tryptophan, 4 µg/ml vitamin B1, 1 mM Mg_2_SO_4_, 0.5 mM CaCl_2_, 0.2% arabinose, and 0 µM and 100 µM FeCl_3_. The growth curve for each culture was based from the A_550_ at various time points (minutes) beginning with the initial absorbance (0 minutes) immediately following the inoculation of the subcultures.

### Proteinase K Accessibility and localization determination

The TonB protein, Maltose-binding protein and the His-tagged peptide proteinase K (PK) accessibility was performed as previously described (49,31,22). Cultures were grown to saturation in Luria-Bertani broth at 37°C with aeration. The saturated cultures were subsequently sub-cultured 1:100 in M9 minimal media supplemented with 0.4% glycerol, 0.2% Casamino Acids, 40 µg/ml tryptophan, 4 µg/ml vitamin B1, 1 mM Mg_2_SO4, 0.5 mM CaCl_2_, and 1.85 µM FeCl_3_. At A_550_ of ∼0.2, cultures were treated with a final concentration of 0.2% arabinose to induce plasmid expression. The whole cell, spheroplast, and lysed spheroplast samples were obtained as previously described (54). The periplasmic fraction samples were obtained from the supernatant of the pelleted spheroplasts. The samples were precipitated with 10% TCA final concentration following a 15 min treatment with PK (25µg/mL) and resuspended in Laemmli sample buffer (55). Equal sample volumes (confirmed from Coomassie staining of the membrane) were loaded onto 16% acrylamide with 6% bis-acrylamide tricine gels and processed for immunoblot analysis (56). The His-tagged peptides, TonB protein, and Maltose-binding proteins were detected by probing with anti-His monoclonal antibodies [EMD Millipore, prod # 70796], anti-TonB 4F1 monoclonal antibodies (57), and anti-Maltose-binding protein (MBP) monoclonal antibodies [New England Biolabs, prod# E8032].

### Ethidium bromide accumulation

Cultures were grown to saturation in Luria-Bertani broth at 37°C with aeration. The saturated cultures were subsequently sub-cultured 1:100 in M9 minimal media supplemented with 0.4% glycerol, 0.2% Casamino Acids, 40 µg/ml tryptophan, 4 µg/ml vitamin B1, 1 mM Mg_2_SO_4_, 0.5 mM CaCl_2_, and 37 µM FeCl_3_. At A_550_ of ∼0.2, cultures were treated with a final concentration of 0.2% arabinose to induce plasmid expression. At *A*_550_ of 0.5, 2 mL of cells were transferred to 4.5 mL disposable cuvettes (VWR Cat# 30620-276) Ethidium bromide (6.25 μM) was added to the cells and fluorescence was detected using a LS-55 fluorescence spectrophotometer. The excitation wavelength was 520 nm and the emission wavelength was 590 nm. The width of the excitation and emission slits were 15 nm. Fluorescence emission data was collected every second for 3 minutes using the Time Drive application from the FLWINLAB software. For CCCP controls, either 15 μM or 60 μM CCCP was added two minutes prior to ethidium bromide. The ethidium bromide accumulation rate for each sample was determined by linear regression and recorded as Fluorescence Units per second (FU/sec).

### *In vivo* formaldehyde-cross-linking

Cultures were grown to saturation in Luria-Bertani broth at 37°C with aeration. The saturated cultures were subsequently sub-cultured 1:100 in M9 minimal media supplemented with 0.4% glycerol, 0.2% Casamino acids, 40 µg/ml tryptophan, 4 µg/ml vitamin B1, 1 mM Mg_2_SO_4_, 0.5 mM CaCl_2_, and 1.85 µM FeCl_3_. At A_550_ of ∼0.2, cultures were treated with a final concentration of 0.2% arabinose to induce plasmid expression. Culture volumes were adjusted to harvest 0.5 *A*_550_ -mL of cells. Bacteria were resuspended in sodium phosphate buffer (pH 6.8) and treated for 15 min at room temperature with 1% formaldehyde (Electron Microscopy Sciences Cat #15710) as previously described (46). Following formaldehyde treatment, bacteria were suspended in Laemmli sample buffer (55), with equal cell numbers loaded in each well of 13% sodium dodecyl sulfate (SDS)-polyacrylamide gels. ExbD complexes and TonB complexes were detected by immunoblotting with ExbD-specific polyclonal antibodies and TonB-specific monoclonal antibodies, respectively (14, 56). Equal loads were confirmed by Coomassie staining of the PVDF membrane used for the immunoblot.

## Supporting information

All supplemental files

