## Supplementary material for "An ExbD Disordered Domain Peptide Inhibits TonB System Activity": All supplemental files

**Kopp, D.R., Jr. and Postle, K.**

### SUPPLEMENTAL INFORMATION

**Table S1: Strains and Plasmids**

| **Strain** | **Genotype** | **Reference** |
|---|---|---|
| W3110 | $F^-$ IN(*rrnD-rrnE*)1 | (1) |
| KP1270 | W3110, *aroB-* | (2) |
| KP1477 | W3110 Δ*tonB*::*kan* | (3) |
| RA1045 | W3110 *ΔexbD, ΔtolQR* | (4) |
| **Plasmid** | **Genotype or Phenotype** | **References** |
| pET-12a | T7-promoter-ompT(ss), $amp^r$, pBR322 ori | (5) |
| pBAD33 | arabinose-inducible, $cam^r$ , p15A ori | (6) |
| pKP1694 | T7 promoter – ompT(ss)-ExbD(44-141) | Present study |
| pKP1695 | T7 promoter – ompT(ss)-TonB(33-239) | Present study |
| pKP1714 | ompT(ss)-ExbD(44-141) | Present study |
| pKP1715 | ompT(ss)-TonB(33-239) | Present study |
| pKP1832 | dsbA(ss)-ExbD(44-141) | Present study |

| | | |
|---|---|---|
| pKP1838 | dsbA(ss)-ExbD(44-91) | Present study |
| pKP1839 | dsbA(ss)-ExbD(92-141) | Present study |
| pKP1847 | dsbA(ss)-ExbD(44-63) | Present study |
| pKP1855 | dsbA(ss)-ExbD(92-121) | Present study |
| pKP1856 | dsbA(ss)-ExbD(122-141) | Present study |
| pKP1946 | dsbA(ss)-ExbD(64-91) | Present study |
| pKP1960 | dsbA(ss)-ExbD(44-63)-His6X | Present study |
| pKP1961 | dsbA(ss)-ExbD(64-91)-His6X | Present study |
| pKP1977 | dsbA(ss)-His6X | Present study |
| pKP1980 | dsbA(ss)-ExbD(92-121)-His6X | Present study |
| pKP1981 | dsbA(ss)-ExbD(54-73)-His6X | Present study |
| pKP1982 | dsbA(ss)-ExbD(82-101)-His6X | Present study |
| pKP1983 | dsbA(ss)-ExbD(112-131)-His6X | Present study |
| pKP2006 | dsbA(ss)-ExbD(122-141)-His6X | Present study |
| pKP2070 | dsbA(ss)-ExbD(44-58)-His6X | Present study |
| pKP2013 | dsbA(ss)-ExbD(44-141)-His6X | Present study |
| pKP2069 | dsbA(ss)-ExbD(49-63)-His6X | Present study |

| | | |
|---|---|---|
| pKP2082 | dsbA(ss)-ExbD(49-63) | Present study |
| pKP2089 | dsbA(ss)-ExbD(44-63; V45A, V47A) | Present study |
| pKP2142 | dsbA(ss)-ExbD(44-63; V45A, V47A) -His6X | Present study |

**ompT(ss)**

Met Arg Ala Lys Leu Leu Gly Ile Val Leu Thr Thr Pro Thr Ala Thr Ser Ser Phe Ala Ser Thr Gly Ser

**dsbA(ss)**

Met Lys Lys Ile Trp Leu Ala Leu Ala Gly Leu Val Leu Ala Phe Ser Ala Ser Ala Ala Gln Tyr Glu

**Figure S1: ompT and dsbA signal peptide and linker sequences.**

The three-letter amino acid codes for the ompT signal sequence [ompT(ss)] (5) and those for dsbA signal sequence [dsbA(ss)] (7) used. The linker sequences are highlighted in gray.

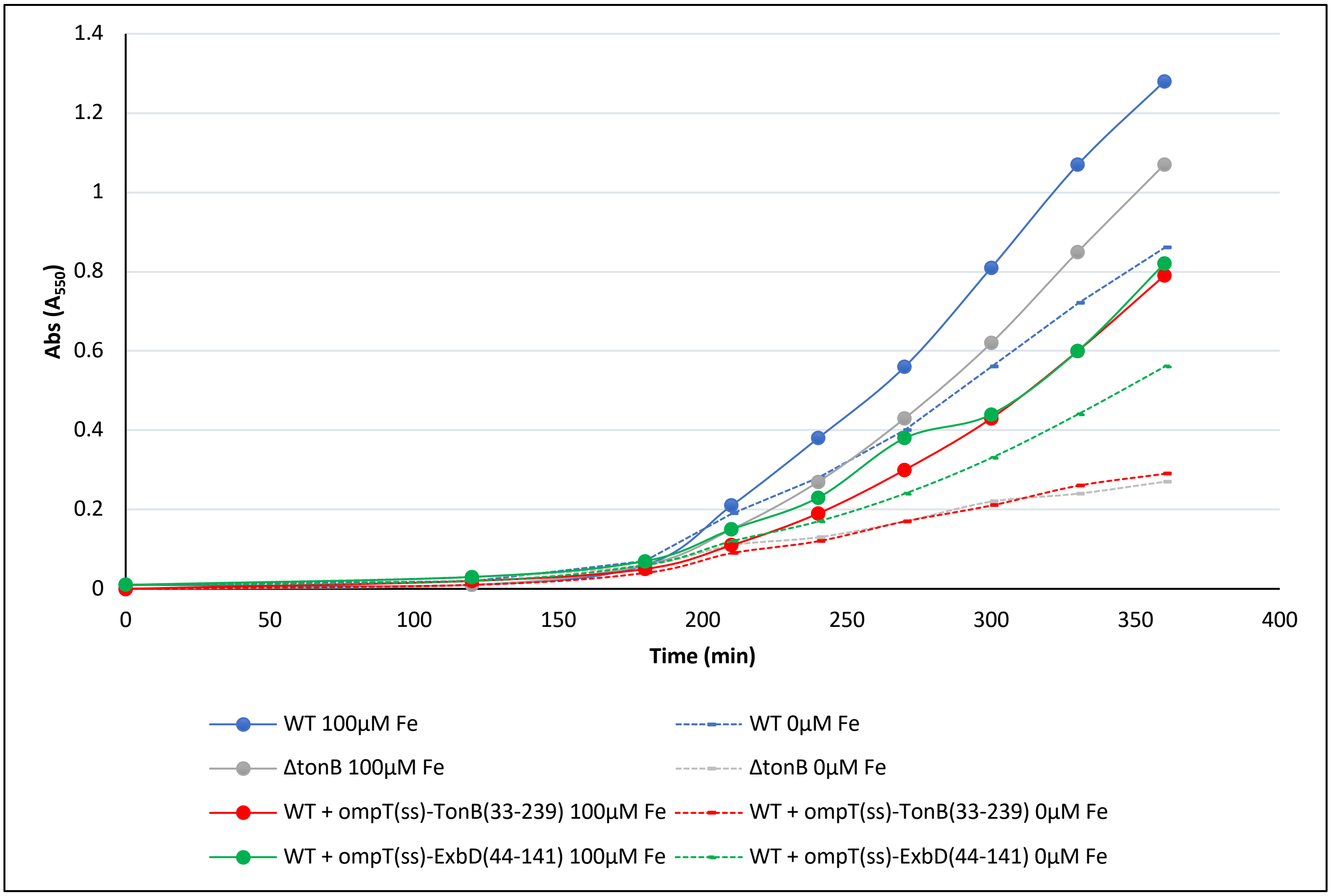


**Figure S2: Cells expressing ompT(ss)-TonB(33-239) have an iron-dependent growth inhibition and Cells expressing ompT(ss)-ExbD(44-141) do not have an iron-dependent growth inhibition.** Overnight cultures were sub-cultured 1:1000 in M9 with 100µM Fe and no added Fe as described in the Materials and Methods and grown at 37˚C with aeration sub-cultured. Arabinose [0.2% (w/v)] was also added to the M9 medium for ompT(ss)-TonB(33-239) and ompT(ss)-ExbD(44-141) expression induction. The growth curve differences between W3110 (WT) expressing either ompT(ss)-TonB(33-239) or W3110 (WT) expressing ompT(ss)-ExbD(44-141) were compared to those of controls W3110 (WT) and KP1477 (ΔtonB). The Y-axis is the absorbance at a wavelength of 550 nM. The X-axis indicates the time following inoculation in minutes.

The solid dots on the growth curves were the absorbances recorded at the specific minutes following inoculation.

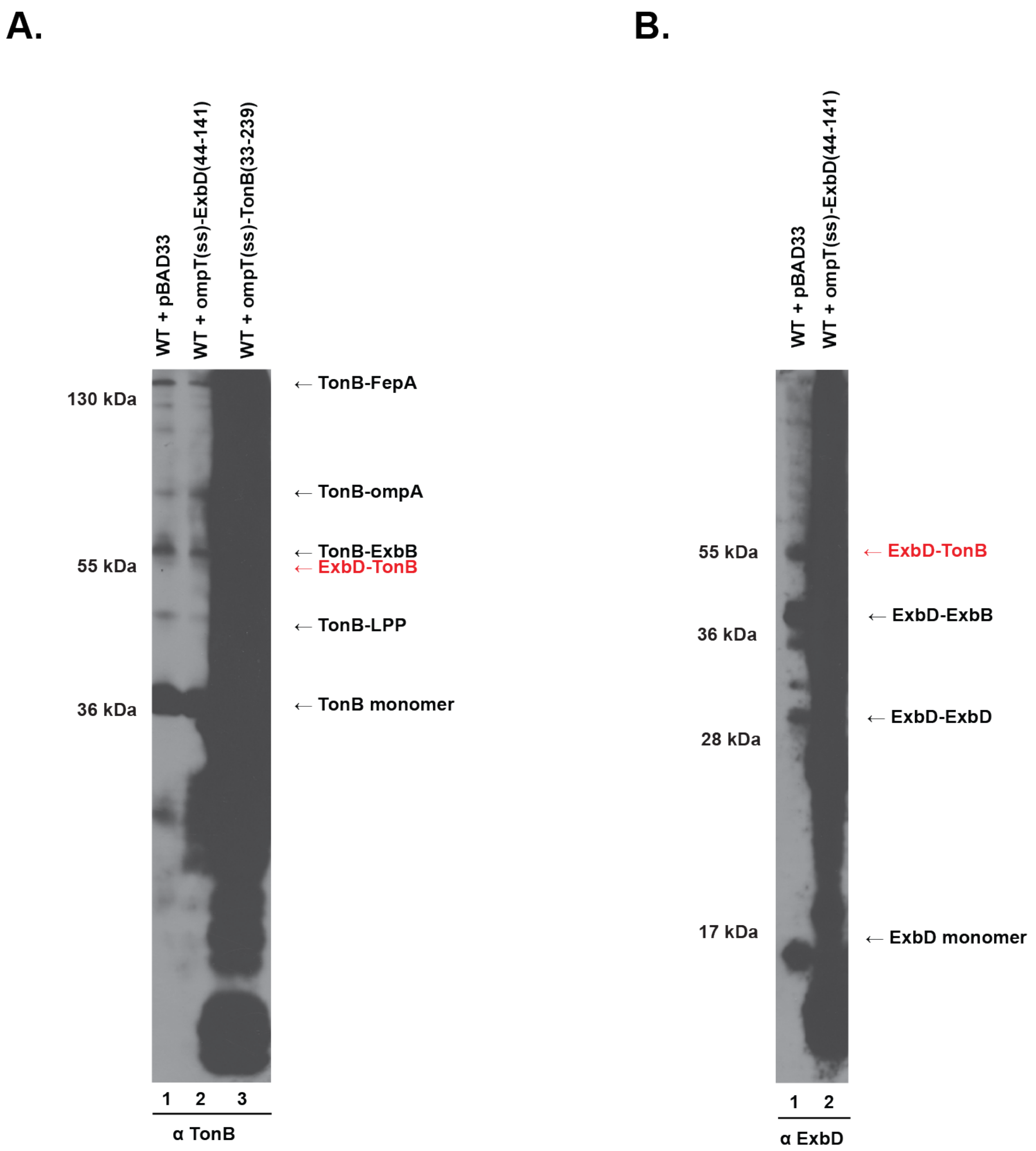


**Figure S3: TonB complexes and ExbD complexes are unable to be identified with expressing, ompT(ss)-TonB(33-239) and ompT(ss)-ExbD(44-141) respectively.**

Cells were grown to mid-exponential phase, at which point formaldehyde-cross-linking was performed as described in Materials and Methods. Equivalent numbers of cells were visualized on immunoblots of 13% SDS-polyacrylamide gels probed with (A) anti-TonB monoclonal antibodies or (B) anti-ExbD polyclonal antibodies. (A, lane 1) WT + pBAD33 refers to W3110 harboring the empty vector pBAD33. (A, lane 2) WT + ompT(ss)-ExbD(44-141) is W3110 expressing plasmid-encoded ompT(ss)-ExbD(44-141). (A, lane 3) WT + ompT(ss)-TonB(33-239) is W3110 expressing plasmid-encoded ompT(ss)-TonB(33-239). Protein expression was induced with 0.2% (w/v) arabinose. (A, right) Positions of the previously characterized TonB formaldehyde cross-linked complexes are shown: TonB-FepA complex, TonB-ompA complex, the TonB-ExbB complex, the PMF-dependent TonB-ExbD complex (red text), TonB-LPP (Braun's lipoprotein) and TonB monomer (9,8). (A, left) Mass markers are shown. (B, lane 1) WT + pBAD33 refers to W3110 harboring the empty vector pBAD33. (B, lane 2) WT + ompT(ss)-ExbD(44-141) is W3110 expressing plasmid-encoded ompT(ss)-ExbD(44-141). (B, right) Positions of the previously characterized ExbD formaldehyde cross-linked complexes are shown: PMF-dependent complex ExbD-TonB complex (red text), ExbD-ExbB heterodimer, ExbD homodimer (ExbD-ExbD) and ExbD monomer (8). (B, left) Mass markers are shown. The plasmid identities are listed in Table S1.

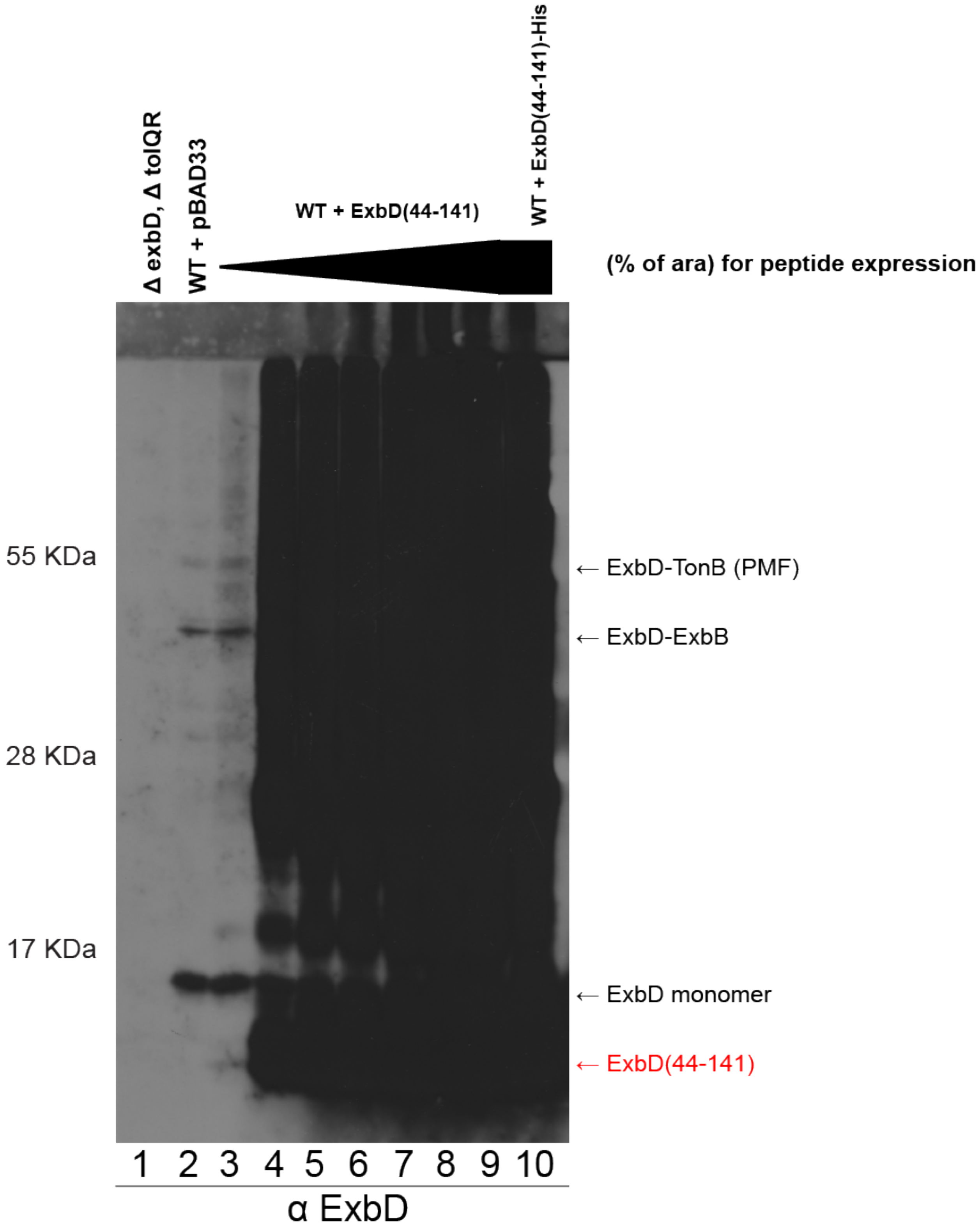
Δ exbD, Δ tolQR
WT + pBAD33
WT + ExbD(44-141)
WT + ExbD(44-141)-His
(% of ara) for peptide expression
55 KDa
28 KDa
17 KDa
← ExbD-TonB (PMF)
← ExbD-ExbB
← ExbD monomer
← ExbD(44-141)
1 2 3 4 5 6 7 8 9 10
α ExbD

**Figure S4: dsbA(ss)-ExbD(44-141) caused too much background to accurately assess its effect on ExbD complexes.** Samples are the same from Fig. 1 probed with anti-ExbD antibodies. Strains were grown to mid-exponential phase, at which point formaldehyde-cross-linking was performed as described in Materials and Methods. Equivalent numbers of cells were visualized on immunoblots of 13% SDS-polyacrylamide gels probed with anti-ExbD polyclonal antibodies. (lane 1) ΔtonB refers to W3110 with the *tonB* gene deleted (KP1477) and (lane 2) WT + pBAD33 refers to W3110 harboring the empty vector pBAD33. (lanes 3-9) WT + ExbD(44-141) is W3110 expressing plasmid-encoded dsbA(ss)-ExbD(44-141) under the arabinose inducible promoter (pKP1832). The expanding black triangle represents the increasing arabinose that was added to each sample: 0%, 0.0025%, 0.005%, 0.01%, 0.02%, 0.05%, and 0.2% (w/v). (lane 10) WT + ExbD(44-141)-His is W3110 expressing plasmid-encoded dsbA(ss)-ExbD(44-141)-His6X under the arabinose inducible promoter (pKP2013) with at 0.2% (w/v) arabinose. (right, labelled in black) Positions of the previously characterized ExbD formaldehyde cross-linked complexes are shown: ExbD-TonB PMF-dependent complex [ExbD-TonB (PMF], ExbD-ExbB, ExbD homodimer (ExbD-ExbD) and ExbD monomer (8). (right, labeled in red) The suspected ExbD(44-141) peptide. (left) Mass markers are shown. The plasmid identities are listed in Table S1.

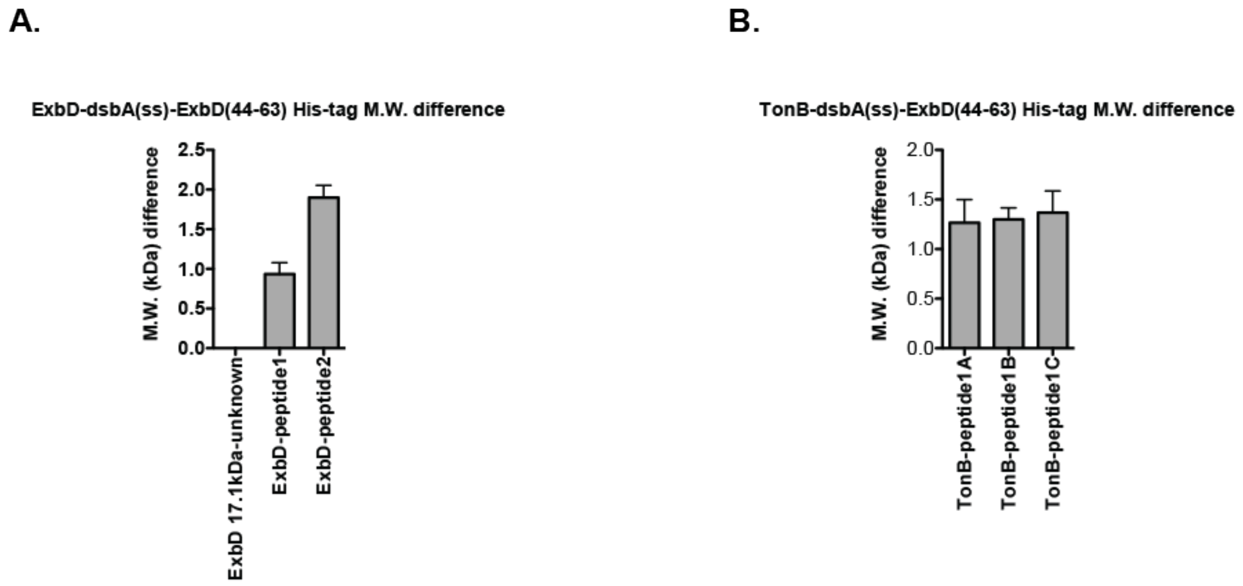


**Figure S5: Molecular weight differences between unknown (A) ExbD and (B) TonB complexes when cells express the dsbA(ss)-ExbD(44-63) peptide and when cells express the dsbA(ss)-ExbD(44-63) peptide dsbA(ss)-ExbD(44-63)-His6X peptide.** A standard curve was made from the distance migrated on the 13% SDS gel of known molecular weights (kDa) from the molecular-weight ladder. The distance of ExbD and TonB unknown complexes were measured and the molecular weights were determined from the standard curve equation calculated from molecular weight ladder. (A) ExbD 17.1 kDa complex weight did not change between dsbA(ss)-ExbD(44-63) peptide and dsbA(ss)-ExbD(44-63)-His6X peptide. ExbD-peptide 1 had a molecular weight increase of ~1 kDa and ExbD-peptide 2 had a molecular weight increase of ~2 kDa when cells expressed dsbA(ss)-ExbD(44-63)-His6X compared to when cells expressed dsbA(ss)-ExbD(44-63). (B) TonB-peptide1A, TonB-peptide1B, and TonB-peptide1C all had a similar molecular weight increase of ~ 1.25 kDa when cells expressed dsbA(ss)-ExbD(44-63)-His6X compared to when cells expressed dsbA(ss)-ExbD(44-63).

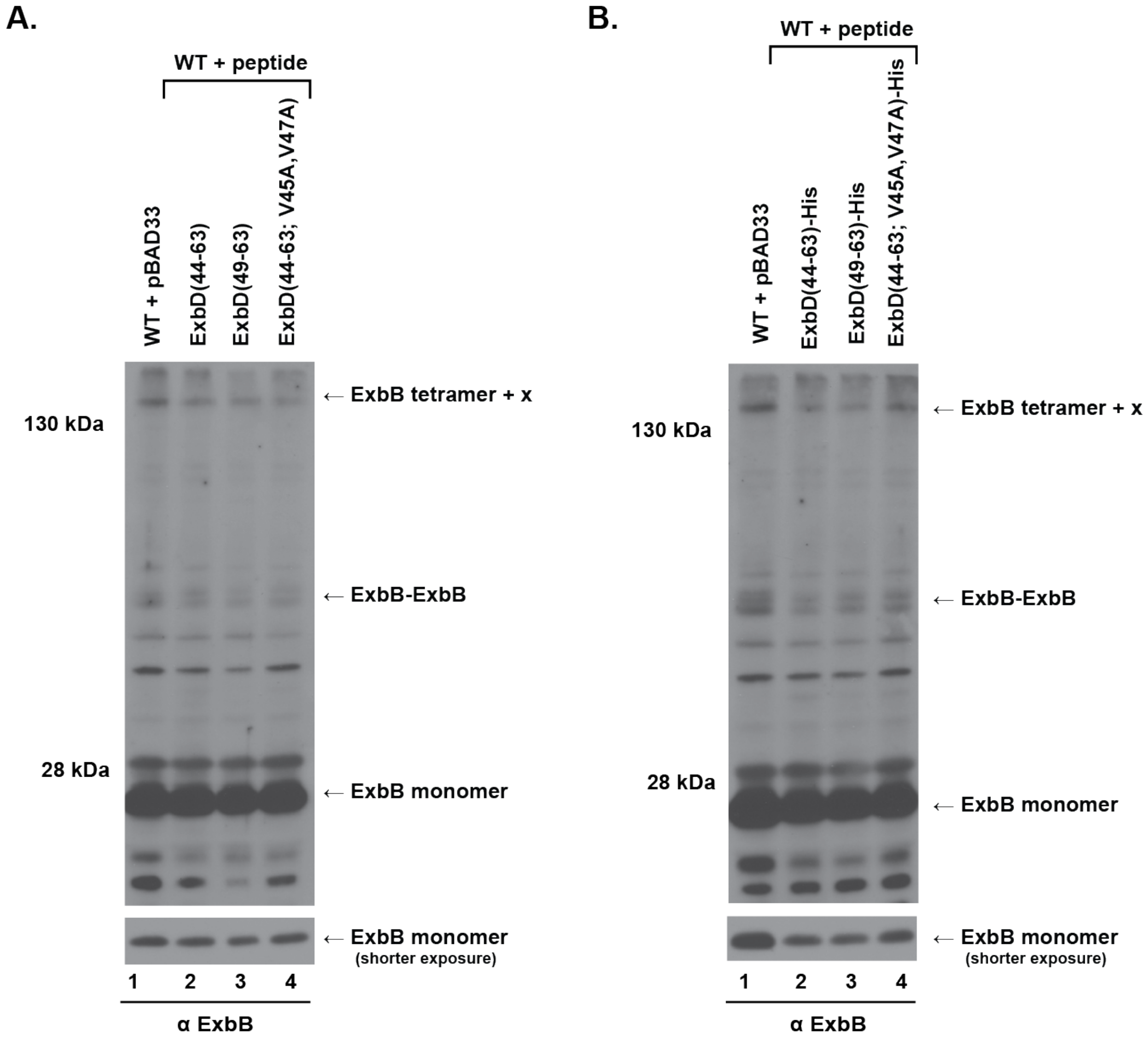


**Figure S6 dsbA(ss)-ExbD(44-63) peptides are not trapped with ExbB**

(A) Formaldehyde cross-linking of W3110 (WT) expressing dsbA(ss)-ExbD(44-63) peptide, dsbA(ss)-ExbD(49-63), or dsbA(ss)-ExbD(44-63; V45A, V47A) probed with anti-ExbB polyclonal antibodies. Cells were grown to mid-exponential phase, at which point formaldehyde-cross-linking was performed as described in Materials and Methods.

Equivalent numbers of cells were visualized on immunoblots of 13% SDS-polyacrylamide gels. (lane 1) WT+ pBAD33 refers to W3110 harboring the empty vector pBAD33. (lane 2) ExbD(44-63) refer to W3110 expressing the plasmid-encoded dsbA(ss)-ExbD(44-63). (lane 3) ExbD(49-63) refer to W3110 expressing the plasmid-encoded dsbA(ss)-ExbD(49-63). (lane 4) ExbD(44-63; V45A, V47A) refer to W3110 expressing the plasmid-encoded dsbA(ss)-ExbD(44-63; V45A, V47A). All plasmids were under the arabinose inducible promoter in the presence of 0.2% (w/v) (A, right) Positions of the previously characterized ExbB formaldehyde cross-linked complexes are shown: the ExbB tetramer complexed with a mystery protein(s), ExbB homodimer (ExbB-ExbB), the ExbB monomer (4). (A, left) Mass markers are shown. (A, bottom) A shorter exposure of ExbB monomer corresponding to each sample. (B) Formaldehyde cross-linking of W3110 (WT) expressing dsbA(ss)-ExbD(44-63)-His6X peptide, dsbA(ss)-ExbD(49-63)-His6X peptide, or dsbA(ss)-ExbD(44-63; V45A, V47A)-His6X peptide probed with anti-ExbB polyclonal antibodies. The formaldehyde cross-linking procedure was the same as in A. (lane 1) WT+ pBAD33 refers to W3110 harboring the empty vector pBAD33. (lane 2) ExbD(44-63)-His refer to W3110 expressing the plasmid-encoded dsbA(ss)-ExbD(44-63). (lane 3) ExbD(49-63)-His refer to W3110 expressing the plasmid-encoded dsbA(ss)-ExbD(49-63)-His6X. (lane 4) ExbD(44-63; V45A, V47A)-His refer to W3110 expressing the plasmid-encoded dsbA(ss)-ExbD(44-63; V45A, V47A)-His6X. All plasmids were under the arabinose inducible promoter in the presence of 0.2% (w/v) arabinose. (B, right) Positions of the previously characterized ExbB formaldehyde cross-linked complexes are shown: the ExbB tetramer complexed with a mystery protein(s), ExbB homodimer (ExbB-ExbB), the ExbB monomer (6). (B, left) Mass markers are shown. (B, bottom) A shorter exposure of ExbB monomer corresponding to each sample. The plasmid identities are listed in Table S1.
